# Direct disassembly of α-syn preformed fibrils into native α-syn monomers by an all-D-peptide

**DOI:** 10.1101/2023.12.11.571053

**Authors:** Marc Sevenich, Ian Gering, Madita Vollmer, Selma Aghabashlou Saisan, Markus Tusche, Tatsiana Kupreichyk, Thomas Pauly, Matthias Stoldt, Wolfgang Hoyer, Antje Willuweit, Janine Kutzsche, Nils-Alexander Lakomek, Luitgard Nagel-Steger, Lothar Gremer, Gültekin Tamgüney, Jeannine Mohrlüder, Dieter Willbold

**Affiliations:** Institute of Biological Information Processing (IBI-7), Forschungszentrum Jülich, Jülich, Germany; Institut für Physikalische Biologie, Heinrich-Heine-Universität Düsseldorf, Düsseldorf, Germany; Priavoid GmbH, Düsseldorf, Germany; Institute of Neuroscience and Medicine (INM-4), Forschungszentrum Jülich, Jülich, Germany; JuStruct, Forschungszentrum Jülich, Jülich, Germany

## Abstract

Parkinson’s disease (PD) is the most common neurodegenerative movement disorder worldwide. One of its central features is the neurodegeneration that starts in the substantia nigra and progressively tends to involve other brain regions. α-Synuclein (α-syn) and its aggregation during pathogenesis have been drawn into the center of attention, where especially soluble oligomeric and fibrillar structures are thought to play a key role in cell-to-cell transmission and induction of toxic effects. Here, we report the development of all-D-enantiomeric peptide ligands that bind monomeric α-syn with high affinity, thereby stabilizing the physiological intrinsically disordered structure and preventing initiation of aggregation, and more important, disassembling already existing aggregates. This “anti prionic” mode of action (MoA) has the advantage over other MoAs that it eliminates the particles responsible for disease propagation directly and independently of the immune system, thereby restoring the physiological monomer. Based on mirror image phage display on the D-enantiomeric full-length α-syn target, we identified SVD-1 and SVD-1a by next generation sequencing, Thioflavin-T screens and rational design. The compounds were analyzed with regard to their anti-aggregation potential and both compounds showed aggregation delaying as well as seed capacity reducing effects in *de novo* and seeded environments, respectively. High affinity towards the monomeric α-syn, in the low nano- to picomolar K_D_ range was identified by surface plasmon resonance (SPR). SVD-1a reduced toxic effects as well as intracellular seeding capacity of α-syn pre-fromed fibrils (PFF) in cell culture. SVD-1a disassembled α-syn PFF into monomers as identified by atomic force microscopy (AFM), time dependent dynamic light scattering (DLS) and size exclusion chromatography (SEC) analysis. The present work provides promising results on the development of lead compounds with this anti-prionic mode of action for treatment of Parkinson’s disease and other synucleinopathies.

## INTRODUCTION

Fibrils consisting of α-syn have been extracted from brain tissue of patients suffering from Parkinson’s disease (PD) . PD, dementia with Lewy bodies (DLB) and multiple system atrophy (MSA) are diseases that are collectively referred to as synucleinopathies, as they are all characterized by the accumulation of insoluble α-synuclein (α-syn) aggregates in neuronal cells. The majority of the filamentous proportion of these deposits is composed of the α-syn protein (*1, 2*). α-Syn, encoded by the *SNCA* gene, is a 140 amino acid, 14.6 kDa, pre-synaptically located, and intrinsically disordered protein (IDP), which is thought to be involved in synaptic vesicle trafficking, synaptic plasticity as well as modulation of neurotransmitter release including dopamine (*3, 4*). More than insoluble α-syn fibrils, smaller soluble versions of them as well as α-syn oligomers are suspected to be responsible for progression of the disease and the spreading of the pathology through the brain.

On-as well as off-pathway oligomers show a clear negative impact on many cellular processes including membrane, proteasome, mitochondria and ER function, as well as inflammation, autophagy, and synaptic transmission (*5–7*). In addition, recent insights suggest that aggregated species of α-syn are able to self-propagate between neuronal cells in a prion-like manner and, therefore, might be the central factor of the progressive nature of disease pathology (*8–10*).

In this sense, the most promising mode of action for PD treatment is the destabilization and direct elimination of the toxic and self-replicating α-syn species. We have therefore termed such a therapeutic strategy and the compounds that realize it, “anti-prionic” (*11*). This term stays in analogy to the term “antibiotic” where elimination of the self-replicating pathological species is also the desired mode of action. In contrast to antibiotics, where it is about killing bacteria by chemical intervention of bacterial enzymes, anti-prionics need to inverse a thermodynamic equilibrium that favors formation of α-syn fibrils and oligomers from α-syn monomers (Fig. 1). This is achieved by compounds that stabilize the IDP conformation of the α-syn monomer.

**Figure 1:**
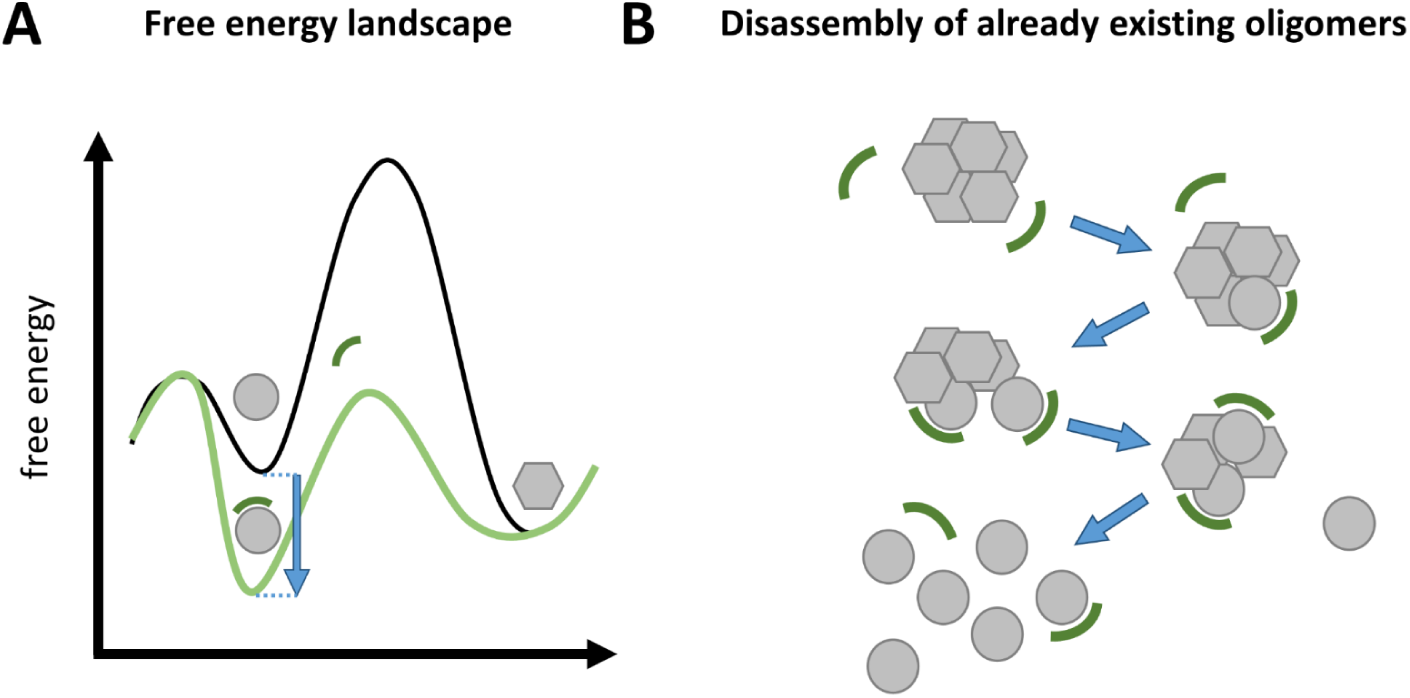
Mechanistic model of the anti-prionic mode of action realized by the all-D-peptide described in the present work. Anti-prionic all-D-peptides are designed to stabilize monomers in their native, intrinsically disordered conformation – symbolized by circles. This conformation is distinct from the yet unknown, but certainly highly defined beta-sheet-rich conformation building blocks in oligomers – symbolized by hexagons. (**A**) Qualitative and schematic free energy landscape for the anti-prionic mode of action. The black line represents the energy landscape in absence of the anti-prionic all-D-peptide. Monomer building blocks in oligomers are more stable than monomers. This allows the formation of oligomers from monomers thermodynamically, although there is a kinetic barrier, which is called primary nucleation and currently under intensive investigation. Stabilization of the monomer by the anti-prionic all-D-peptide is lowering the free energy of the monomer (light green line), when in complex with the all-D-peptide, by the free binding energy (blue arrow) of the complex. Because in the presence of the anti-prionic all-D-peptide, the monomer has a lower free energy as compared to the oligomer, oligomers are disassembled into monomers. (**B**) Mechanistic model for disassembly of already existing oligomers from top to bottom: Anti-prionic all-D-peptides – symbolized by circle segments – approach oligomers. Due to their affinity to α-syn monomers, each all-D-peptide will interact with one of the α-syn building blocks within the oligomer assembly and thereby pushes its conformation towards the intrinsically disordered monomer conformation. This is incompatible with the oligomer assembly and therefore destabilizing the oligomer assembly. Further destabilization by interaction of additional anti-prionic molecules with other monomer building blocks, ultimately leads to the complete disassembly of the oligomer into monomers in their native intrinsically disordered conformation. Both molecules remain disordered in this transient complex, which may therefore be called “fuzzy complex” (*12*). We called this mode of action “anti-prionic”, because it is ultimately disrupting prion-like behaving aggregates (*11*).

The success of this therapeutic strategy against the target Aβ oligomers for the treatment of Alzheimer’s disease (AD) was previously demonstrated with the anti-prionic compound RD2 (*13–15*). RD2 is a 12-aa all-D-enantiomeric peptide that binds Aβ monomers with high affinity (*16*). This results in the destabilization and ultimately the disassembly of oligomers into monomers. This kind of target engagement has been demonstrated *in vitro* (*14*), *in vivo* (*13*) and most recently, *ex vivo* with patient brain derived Aβ oligomers (*17*). Moreover, the compound proofed cognitive restoration when administered to AD-mice with full-blown pathology (*13*), while its D-enantiomeric structure and low molecular weight confers proteolytic stability, high bioavailability and penetration of the blood-brain barrier (*14, 18, 19*). In this study, we demonstrate the realization of the anti-prionic MoA for α-syn. The developed all-D-enantiomeric peptides SVD-1 and SVD-1a bind monomeric α-syn with high affinity, inhibit non-seeded as well as seeded α-syn aggregation, and most importantly disassemble pre-formed fibrils (PFF) into monomers with high efficiency.

## MATERIAL AND METHODS

### Recombinant expression and purification of monomeric wt and α-syn A140C

N-terminal acetylated α-syn wt (hereinafter referred to as α-syn) and acetylated α-syn-A140C were expressed in *E. coli* BL21(DE3) carrying codon-optimized α-syn in pT7 vector and the pNatB vector with the N-terminal acetylation enzyme from *Schizosaccharomyces pombe* (*20*). Expression was performed in LB or ^15^N-supplemented M9-minimal medium with 1 mM IPTG after reaching an OD_600_ of 1.2 followed by incubation for 4 h at 37 °C. Purification was performed as described previously (*21*) with some modifications: The pellets from 1 l expression was resuspended in 25 ml 20 mM Tris pH 8.0 and boiled at 95 - 100 °C for 2 x 15 min. After centrifugation at 20.000 x g for 30 min at 4 °C, the supernatant was precipitated using a final concentration of 0.45 g/ml of ammonium sulfate. The protein was pelleted at 20.000 x g for 30 min and resuspended in 50 ml 20 mM Tris-HCl pH 8.0. After sterile filtration the sample was loaded on a HiPrep QFF 16/10 (Cytiva, USA, CV = 20 ml) anion exchange column. Gradient elution was performed with a target concentration of 800 mM NaCl over 20 CV. Recombinant α-syn eluted at a conductivity of 28-32 mS/cm. The fractions containing recombinant α-syn were pooled and precipitated using ammonium sulfate as described previously. The pellets were resuspended in 5 ml 50 mM Tris-HCl pH 7.4 50 mM NaCl and loaded on a HiLoad Superdex 60/75 pg gel filtration column (Cytiva, USA, CV = 120 ml). The expression yielded 20-30 mg/l as determined by A_275_ with an extinction coefficient of 5600 M^-1^ cm^-1^. Protein aliquots were frozen with liquid nitrogen and stored at -80 °C.

### Mirror-image phage display selection

In phage display, exogenous peptides are presented on phage particles by fusion with the major coating proteins. Consecutive rounds of biopanning and amplification increase the fraction of phages presenting strong target binders, which is detectable by sequencing of the variable portion of the genome (*22*). Using mirror-image phage display, the L-enantiomeric selection target is replaced by an otherwise identical D-enantiomeric version. This allows the identification of D-enantiomeric peptides that show high affinity for the physiological L-target and are more resistant to metabolic degradation than their L-enantiomeric counterparts (*23, 24*). For mirror-image phage display, the commercially available M13-bacteriophage library TriCo-16 (Creative Biolabs, USA) was used. The library has a capacity of 2.6 · 10^10^ pIII fused 16-mer peptide variants. D-enantiomeric full-length α-syn carrying a C-terminally biotinylation and N-terminal acetylation was purchased as lyophilized powder from P&E (Peptides and Elephants, GE) with a purity of > 90 %. To minimize non-target related peptide enrichment, the display format was alternated between a polystyrene and polypropylene streptavidin functionalized 96-well plate surface (Maximum capacity plates, BioTeZ, GE). For target immobilization, D-enantiomeric α-syn was diluted to a concentration of 2 pmol/well. Non-coupled streptavidin was quenched using biotin. The selection was performed as described previously (*25–27*) with some minor modifications. Briefly, three consecutive selection rounds were performed using alternating blocking conditions with bovine serum albumin (BSA) and milk powder (MP) with PBS pH 7.4 as selection buffer. Selection pressure was stepwise increased with each selection round using 5 to 10 washing repetitions. The selection was performed in three consecutive rounds on the target (*target selection* = TS). Additionally, input phages resulting from selection rounds on the target were incubated on a surface without target (*direct control* = DC), which was otherwise treated identically to the TS surface. As a second control, a consecutive selection was performed exclusively without target on otherwise identically treated surfaces (*empty selection* = ES). A concentration of 2 · 10 ^12^ CFU ml^-1^ was used as input for all selection rounds and controls.

### Enrichment ELISA

Enrichment ELISA is a method to identify enrichment of target binding phages during phage display selection. All steps were performed as described previously with some minor changes (*25*). Briefly, 20 pmol/well of the D-enantiomeric α-syn target was immobilized on a streptavidin coated polystyrene 96-well plate (maximum capacity plates, BioTeZ, GE). Both, the target immobilized and target free surface were quenched with biotin In total, 2.5 · 10^11^ phages from the TS input samples were diluted in 100 µl washing buffer and incubated on target and control surface. The A_450_ of the product of the peroxidase reaction product 3,3’,5,5’-tetramethylbenzidine diimine was quantified after reaction stop with H_2_SO_4_ by absorption measurement in a Fluorostar optima platereader (BMG labtech, GE; n = 3).

### ssDNA purification and next generation sequencing of phage input samples

The ssDNA of the input phage suspensions resulting from mirror-image phage display selection was purified by phage precipitation and subsequent ssDNA separation as described previously (*25, 28, 29*). For next generation sequencing, PCR was performed adding adapter sequences to both, the 3’ and 5’ end of the amplification product. Amplicon next generation sequencing was performed by BMFZ-GTL Düsseldorf (GE) with a MiSeq system (Illumina, USA).

### Analysis of the next generation sequences by filtering and clustering

Variable DNA sequences resulting from next generation sequencing (NGS) were transcribed to peptide sequences as described previously (*25, 28, 29*). The transcribed sequences were filtered based on their frequency increase in TS (library <TS1 < TS2 < TS3), their correlation with the presence of the target (TS2 > DC2; TS3 > DC3) and their frequency in the selection without target (TS1 > ES1; TS2 > ES2; TS3 > ES3). Sequences that passed the filter were ranked corresponding to their enrichment from library to TS3 (TS3/library = *enrichment score*) and their frequency in TS3 compared to ES3 (TS3/ES3 = *empty score*). Filtered sequences were used as input for *Hammock clustering software* (*30*). The input FASTA-file included all filtered sequences as ranked by their *empty score*. *Hammock clustering software* was run in full mode with sequences ranked according to their input position (-R input). Cluster motif logos were generated from initial clusters after greedy clustering using the *WebLogo 3.4* application.

### D-enantiomeric peptides

D-enantiomeric peptides were purchased with C-terminal amidation from CASLO (CASLO, DK) as lyophilized chloride salt powder with a purity of > 95 %. SVD-1 and SVD-1a were tested in different buffer conditions including PBS pH 7.4, where UV-vis absorption measurements after 1 h incubation at 37 °C and 20.800 xg centrifugation showed that both compounds completely retained in the supernatant up to at least 1 mM initial concentration.

### Thioflavin T assay

The Thioflavin T (ThT) assay is commonly used for the visualization of α-syn fibrilization, since the dye ThT is able to bind to the amyloidogenic cross-β-sheet proportions of fibril structures. Recombinant α-syn was thawed on ice and centrifuged for 30 min at 21.000 x g and 4°C. The concentration of the supernatant was determined as described previously. Lyophilized D-peptides were thawed at RT for 1 h and dissolved in 500 µl PBS pH 7.4. After centrifugation for 30 min at 21.000 x g the supernatant concentration was determined by UV-vis using the corresponding extinction coefficient at A_280_. All ThT assay experiments were performed at 37°C with 15 µM ThT and 0.05 % sodium azide (w/v) in PBS pH 7.4 if not otherwise stated. ThT fluorescence was monitored with bottom optics at λex = 448 nm and λem = 482 nm in a fluorescence plate reader with orbital averaging on 3 mm (Clariostar or Polarstar Optima, BMG labtech, GE). Prior to measurement start, 120 µl sample solution was transferred to a non-binding 96-half area well plate with transparent flat bottom (Corning, USA). One borosilicate glass bead (d = 3.0 mm, Hilgenberg, GE) was added to each well for all *de novo* aggregation assays. For seeded aggregation assays no bead was used and samples were incubated under quiescent conditions. Wells that surrounded the sample wells were filled with the same volume of buffer to improve heat distribution. Experiments were performed with five replicates (n = 5) if not otherwise stated.

ThT assay was used for different purposes. First, *de novo* ThT aggregation assays served as screening platform for the aggregation delay with the synthetic D-peptides. Here, 50 µM recombinant α-syn was incubated with a three-fold molar excess of each D-peptide, respectively. Samples were shaken before each cycle using orbital shaking mode at 300 rpm for 30 s. Peptides that were insoluble in aqueous buffer were dissolved in 2.5 µl DMSO (0.5 mg peptide) and gradually mixed with PBS pH 7.4 until a final concentration of 2.5 % (v/v) DMSO was reached. For these samples, the reference aggregation of α-syn alone was also performed in PBS pH 7.4 with 2.5 % (v/v) DMSO. Aggregation curves of each replicate were individually fitted using a symmetric Boltzmann sigmoidal fit (OriginPro 2020, OriginLab, USA) with the following formula: 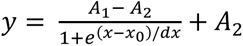 (A_1_ = initial value, A_2_ = final value, x_0_ = inflection point [s], dx = time constant [1/s]). The inflection point of the fit determines the aggregation half-time t ½, whereas the lag-time was approximated with the following formula: t_1/2_ – 2 · dt_1/2_, where dt_1/2_ is defined as the slope of the fit at x = t_1/2_ in 1/s (*31*). Half-time and lag-time were calculated as mean value of the separate fits. For the concentration dependency of aggregation delay, different compound concentrations were applied to 50 µM recombinant α-syn. Lag-time and t ½ were calculated as described previously. For graphical representation, the fitted steady-states were normalized to 1 for all conditions and the mean error was calculated based on the fits. Statistical testing on the significance of the time shifts was performed using the two sample Welch‘s t-test with p < 0.05 (OriginPro 2020, OriginLab, USA). For *de novo* ThT assays including sub-stoichiometric compound concentrations, 10 µM α-syn was used. In contrast to the screening aggregation assay α-syn samples were continuously shaken at 300 rpm using orbital shaking mode to reduce the aggregation time taking measurements every 5 min. Statistical evaluation was performed as described previously by comparing the significance of t ½, lag-time shifts and steady state reduction in the presence of the inhibitor and the same concentration of the corresponding control peptide. For seeded ThT assays, 50 nM monomer equivalent PFF oligomers were incubated ON at 37 °C under quiescent condition together with or without different concentrations of the compounds in the fluorescence plate reader. After 20 h 20 µM monomeric α-syn was added to the samples mixture to start seeding. Measurements were taken every five minutes (n = 3).

### Preparation of PFF oligomers

PFF oligomers were prepared as described previously with some modifications (*32*). First, insoluble PFF were produced by incubation of 300 µM recombinant α-syn in a LoBind reaction tube (Eppendorf GmbH, GE) with one borosilicate glass bead (d = 3.0 mm; Hilgenberg, DE) in 20 mM NaPi pH 7.0 150 mM NaCl 0.05 % (w/v) sodium azide for one week at 37 °C. The insoluble PFF were harvested by ultracentrifugation at 100.000 x g for 30 min at 4 °C and the pellet was washed several times with 20 mM NaPi pH 7.0 150 mM NaCl. The monomer equivalent concentration was determined by measuring the α-syn concentration in the supernatant after the first centrifugation and subtracting it from the start concentration for fibrilization. The insoluble PFF were resuspended in buffer and frozen at -80 °C with liquid N_2_. PFF oligomers were generated by harsh sonication of 200 µl insoluble PFF with 300 µM monomer equivalent concentration for 3 x 15 s (1 sec. on/off) and 60 % amplitude with a tip sonifier (MS 72 micro tip, Sonopolus, Brandelin, GE). Insoluble PFF were separated by centrifugation at 100.000 x g for 1 h at 4 °C. The supernatant containing PFF oligomers was separated, aliquoted and frozen at - 80 °C with liquid N_2_.

### Surface plasmon resonance kinetic experiments

Measurements were performed using an 8K Biacore device (Cytiva, USA). Interactions were measured using single cycle kinetics experiments. For all assays, the peptide compounds were immobilized as ligand on the sensor surface and recombinant α-syn was injected as analyte in the flow. SVD-1 and SVD-1a were immobilized via primary amino groups on a CMD200M carboxyldextran matrix chip (Xantec, GE). Immobilization was performed after 7 min EDC/NHS activation at 10 µl/min with 50 µg/ml peptide in 10 mM NaAc pH 5.0 for SVD-1 and pH 7.0 for SVD-1a until a saturation signal was reached (SVD-1: 400 RU, SVD-1a: 500 RU). Surface quenching was performed using 1 M ethanolamine pH 8.3. The kinetic experiments were performed using a flow rate of 30 µl/min in PBS pH 7.4 if not otherwise stated. The surface was regenerated in between cycles using 30 s injections of 2 M Gua-HCl at 30 µl/min. Data evaluation was performed using Biacore *insight evaluation software* v3.0 (Cytiva, USA).

### SVD-1a_Cys_MTSL spin label preparation

The spin-labeled analogue of SVD-1a was prepared by covalent attachment of MTSL (S-(1-oxyl-2,2,5,5-tetramethyl-2,5-dihydro-1H-pyrrol-3-yl)methyl methanesulfonothioate, Toronto Research Chemicals, USA) to the C-terminal D-cysteine residue of SVD-1a_Cys. MTSL was dissolved in DMF (N,N-dimethylformamide) with a concentration of 20 mM and diluted in 200 mM HEPES pH 7.6 to a final concentration of 2 mM. 900 µl of the solution was then added to 1 mg of lyophilized SVD-1a_Cys to create a fivefold molar excess of MTSL compared to SVD-1a_Cys. The reaction mixture was incubated for 2 at RT and subsequently applied to a semipreparative RP-HPLC C8 column (Zorbax-300 SB, Agilent, GE) connected to an HPLC system (Agilent 1260, Agilent, GE). Purification of the spin-labeled peptide SVD-1a_Cys_MTSL was achieved by applying an aqueous acetonitrile (ACN) gradient (8% ACN, 0.1% trifluoroacetic acid (TFA) to 60% ACN, 0.1% TFA in Milli-Q water within 40 min), running at a flow rate of 4 ml min^-1^ at 25 °C with a detection at 214 nm. The purified reaction product was flash-frozen with liquid N_2_ and lyophilized (LT-105, Martin Christ, GE). The purity of the SVD-1a_Cys_MTSL spin labeled peptide was verified by RP-HPLC with > 98 %.

### NMR spectroscopy

Samples were prepared at final concentrations of 25 µM ^15^N-labeled full-length acetyl-α-syn in the absence (reference) and presence of an equimolar amount of SVD-1a (not isotopically labeled, thus NMR-invisible) in PBS buffer, pH 7.4, with an addition of 5 % D_2_O for internal reference. 2D ^1^H-^15^N HSQC spectra were recorded back-to back, on a Bruker AVANCE NEO spectrometer (Bruker, USA) operating at 1200 MHz proton Larmor frequency. The experimental temperature was 10 °C. Spectral dimensions were 16.02 ppm (^1^H) x 30 ppm (^15^N), with 2048 points in the ^1^H dimension and 256 increments in the ^15^N dimension, resulting in an acquisition time of 53 ms for the ^1^H dimension and 35 ms for the ^15^N dimension. For each increment, 32 scans were recorded, with a recovery delay of 1 s between scans, resulting in an overall experimental time of 4.8 h per spectrum.

NMR Paramagnetic Relaxation Enhancement (PRE) data were recorded on 25 µM ^15^N-labeled full-length acetyl-α-syn in presence of 25 µM paramagnetically labeled (but not isotopically enriched) SVD-1a_Cys_MTSL (using a MTSL spin-label covalently attached to the SVD-1a C-terminus), resulting in a 1:1 ratio of α-syn : SVD-1a. Intensities (I_para_) were extracted from 2D ^1^H-^15^N Best-TROSY NMR spectra recorded at 600 MHz and 10 °C, each with 128 scans per increment, resulting in a total experimental time of 16 h per spectrum. Reference data were obtained by adding a 20-fold molar excess of ascorbic acid to the same sample, therefore quenching the paramagnetic effect of the spin-label and obtaining a diamagnetic reference sample. The diamagnetic reference spectra and intensities (I_dia_) were recorded back-to-back and under identical conditions as for the paramagnetic sample.

NMR data sets were processed using the Bruker TopSpin software (version 4.1.1) and visualised using *CcpNmr Analysis* (v2.4.2) (*33*). For the assessment of chemical shift changes of α-syn resonances in presence of SVD-1a relative to the reference spectrum (without SVD-1a), peak positions were extracted using the *CcpNmr Analysis* software. From the residue specific chemical shift changes in the ^1^H and ^15^N dimensions, an absolute chemical shift change, Δδ was calculated using the formula 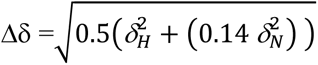 (*34*). For analysis of the PRE data, resonance intensities of the paramagnetic sample and the diamagnetic reference sample peak intensities were extracted using *CcpNmr Analysis*.

### Size exclusion chromatography (SEC)

Elimination of PFF oligomers was evaluated by SEC and subsequent detection of the PFF oligomer and monomer peaks. PFF oligomers were incubated with or without SVD-1a in PBS pH 7.4 for 3 d at 37 °C under quiescent condition in low retention Eppendorf tubes (Eppendorf, GE). Prior to injection, samples were centrifuged at 20.800 xg for 2 min and 100 µl sample was injected on a SEC-HPLC column (Bio SEC-3 300 Å, Agilent, USA) using an Agilent 1260 Infinity II system (Agilent, USA) with a flow rate of 1 ml/min and PBS pH 7.4 as mobile phase. Protein was detected using A_214_.

### Dynamic light scattering (DLS)

Measurements were performed using a SpectroSize 300 131 (XtalConcepts, GE) instrument and a sample volume of 1 ml in a sealed quartz cuvette (Hellma Group, GE). Samples were incubated at 37 °C under quiescent conditions. Prior to measurements, all samples were centrifuged at 21.000 × g for 30 min at 4 °C in order to remove potential impurities from solution. For time dependent DLS measurements data points were recorded every 30 s. Diffusion coefficients were obtained from analysis of the decay of the scattered intensity autocorrelation function and were used to determine apparent hydrodynamic radii via the Stokes-Einstein equation.

### Atomic forced microscopy (AFM)

Samples were prepared by dilution to 1 µM α-syn monomer concentration and 5 µl was incubated and dried on a freshly cleaved mica surface. Surfaces were then three-times washed with 200 µl ddH_2_O and dried using a gentle stream of N_2_. Measurements were performed in a Nanowizard 3 system (JPK BioAFM - Bruker Nano GmbH, GE) using intermittent contact mode with 2 x 2 and 5 x 5 µm section and line rates of 0.5–2 Hz in ambient conditions using a silicon cantilever and tip with nominal spring constant of 26 N/m, average tip radius of 9±2 nm and a resonance frequency of approximately 300 kHz (Olympus OMCL-AC160TS-R3). The images were processed using JPK data processing software (version spm-5.0.84). For the height profiles presented, a polynomial fit was subtracted from each scan line, first independently and then using limited data range.

### Circular dichroism (CD) spectroscopy

Far-UV circular dichroism (CD) data were collected using a Jasco J-1100 spectropolarimeter (Jasco, GE). 350 µl samples were pooled and loaded into a high precision quartz cuvette with a path length of 1 mm (Hellma group, GE). A scan speed of 20 nm/min with five accumulations per sample was performed using far UV wavelengths from 260 to 190 nm. Baseline was corrected by subtracting measurements of the buffer only.

### Cell assay for α-synuclein aggregation

A construct encoding full-length A53T-mutated human α-syn fused with YFP at the C-terminus was synthesized and introduced into the pMK–RQ expression vector (GeneArt; Thermo Fisher Scientific, USA). The α-synA53T–YFP construct was subcloned into the pIRESpuro3 vector (Clontech; Takara Bio, JPN) using NheI (5′) and NotI (3′) restriction sites. HEK293T cells (American Type Culture Collection) were cultured in high glucose Dulbecco’s Modified Eagle’s Medium (DMEM; Sigma-Aldrich, USA) supplemented with 10 % fetal calf serum (Sigma-Aldrich, USA), and 50 units/ml penicillin as well as 50 μg/ml streptomycin (Sigma-Aldrich, USA). Cells were cultured in a humidified atmosphere of 5 % CO_2_ at 37 °C. Cells plated in DMEM were transfected using Lipofectamine 2000 (Invitrogen; Thermo Fisher Scientific, USA). Stable cells were selected in DMEM containing 1 μg/ml puromycin (EMD Millipore, USA). Monoclonal lines were generated by fluorescence-activated cell sorting of a polyclonal cell population in 96-well plates using a MoFlo XDP cell sorter (Beckman Coulter, USA). Finally, the clonal cell line B5 was selected from among 24 clonal cell lines and is referred to as αSynA53T–YFP cells. Peptides were incubated with 1.5 % Lipofectamine 2000 in OptiMEM for 2 h at room temperature. The α-synA53T–YFP cells were plated in a 384-well plate with poly-D-lysine coating (Greiner, AT) at a density of 1,000 cells per well with 0.1 μg/ml Hoechst 33342 (Thermo Fisher Scientific, USA) and the previously prepared transfection mix was added directly to the cells in the well. To seed cellular aggregation of α-syn in α-synA53T–YFP cells, 30 nM soluble α-syn PFF oligomers were incubated with 1.5 % Lipofectamine in OptiMEM for 2 h at room temperature and added to each well 3 h after the first transfection. The plate was then incubated in a humidified atmosphere of 5 % CO_2_ at 37 °C. On day 3 the cells were imaged with an IN Cell Analyzer 6500HS System (Cytiva, USA) using the blue and green fluorescence channel, and analyzed using IN Carta Image Analysis Software (Cytiva, USA) after an algorithm was established to identify intracellular aggregates in living cells. For each condition we used four wells and took 16 images per well, which were analyzed by a fully automated algorithm to avoid bias. Statistical analysis was performed using one-way ANOVA followed by Dunnett’s multiple comparisons test (GraphPad Prism 9, GraphPad Software, USA). Error bars represent standard deviation.

### Cell-viability assay (CellGlo test)

We used the CellTiter-Glo Luminescent Cell Viability Assay (Promega GmbH, GE) to determine the number of viable cells in culture based on quantitation of the ATP present, an indicator of metabolically active cells. After culturing cells in 384-well plates for three days, 35 µl of medium was removed from the wells and 40 µl of CellTiter-Glo Reagent directly added to each well. After mixing, luminescence was measured 10 min later using a Fluostar (BMG labtech, GE).

### Immunofluorescent cell staining

After culturing cells for 3 days on 384-well plates, the cells were fixed in 4 % formaldehyde (Sigma-Aldrich, USA) in PBS (pH 7.4) for 15 min. After washing three times with PBS for 5 min each, the cells were permeabilized with 0.25 % Triton X-100 (Sigma-Aldrich, USA) in PBS for 10 min. After another three washes with PBS for 5 min each, the cells were blocked with 1 % bovine serum albumin (Sigma-Aldrich, USA) in PBS supplemented with 0.1 % Tween 20 (Sigma-Aldrich, USA) for 30 min. The cells were stained with CF633 (Biotium, USA) fluorescently labeled antibodies at 8 µg/ml in 1 % bovine serum albumin in PBS supplemented with 0.1 % Tween-20 for 1–3 h at room temperature in the dark. For detecting total α-syn, we used the anti-α-syn antibody syn211 (Abcam, UK). For detecting oligomeric and fibrillar α-syn, we used the anti-aggregated α-syn antibody, clone 5G4 (Sigma-Aldrich, USA). For detecting α-syn phosphorylated at serine 129, we used the recombinant anti-α-syn (phospho S129) antibody EP1536Y (Abcam, UK). After a final three washes in PBS for 5 min each, the cells were imaged in PBS using an IN-Cell Analyzer 6500HS System and 40-fold magnification (Cytiva, USA).

### Cell viability assay (MTT test)

The potential cell viability rescue of PC12 cells (Leibniz Institute DSMZ, GE) from α-syn toxicity through addition of SVD-1, SVD-1a or SVD-1_scrambled was measured in a MTT (3-(4,5-dimethylthiazol-2-yl)-2,5-diphenyltetrazolium bromide) cell viability test. PC12 cells (Leibniz Institute DSMZ, GE) were cultivated on collagen A-coated (Biochrom GmbH, GE) tissue culture flasks in RPMI 1640 medium supplemented with 5 % fetal calf serum and 10 % horse serum in a 95 % humidified atmosphere with 5 % CO_2_ at 37 °C. 10.000 cells per well in a volume of 100 µl were seeded on collagen A-coated 96-well plates (Thermo Fisher Scientific, USA) and were incubated for 24 h at 37 °C and 300 rpm in a thermo cycler. Then, final concentrations of 30 nM α-syn either in the absence or after pre-incubation with 15 µM SVD-1, SVD-1_scrambled or 0.5 µM SVD-1a was added to the cells. In addition, 15 µM of the peptides alone, cell media, buffer without peptides and 0.1 % Triton X-100 (cytotoxic compound) served as controls. After further incubation in a 95 % humidified atmosphere with 5 % CO_2_ at 37 °C for 24 h, cell viability was measured using the Cell Proliferation Kit I (MTT) (Roche Applied Science, CH) according to manufacturer’s protocol. The MTT formazan product was quantified by measuring the absorbance at 570 nm corrected by subtraction of the absorbance at 660 nm in a FluoroStar Optima plate reader (BMG labtech, GE). All results were normalized to untreated cells grown in medium only. Test on significance was performed using one-way ANOVA with Bonferroni post hoc analysis (OriginPro 2020, OriginLab, USA; n = 4).

## RESULTS

A phage display selection against the mirror image of full-length α-syn yielded SVD-1 as the most promising all-D-peptide for stabilization of native α-syn monomers (for full description of the phage display and peptide screening procedure please see SI Fig. 1 to 6). SVD-1a is a derivative of SVD-1, with D-methionine replaced by D-leucine at position 2, D-lysine replaced by D-arginine at position 1, and addition of five D-arginines to the C-terminal end yielding the sequence shown in fig. 2.

**Figure 2:**
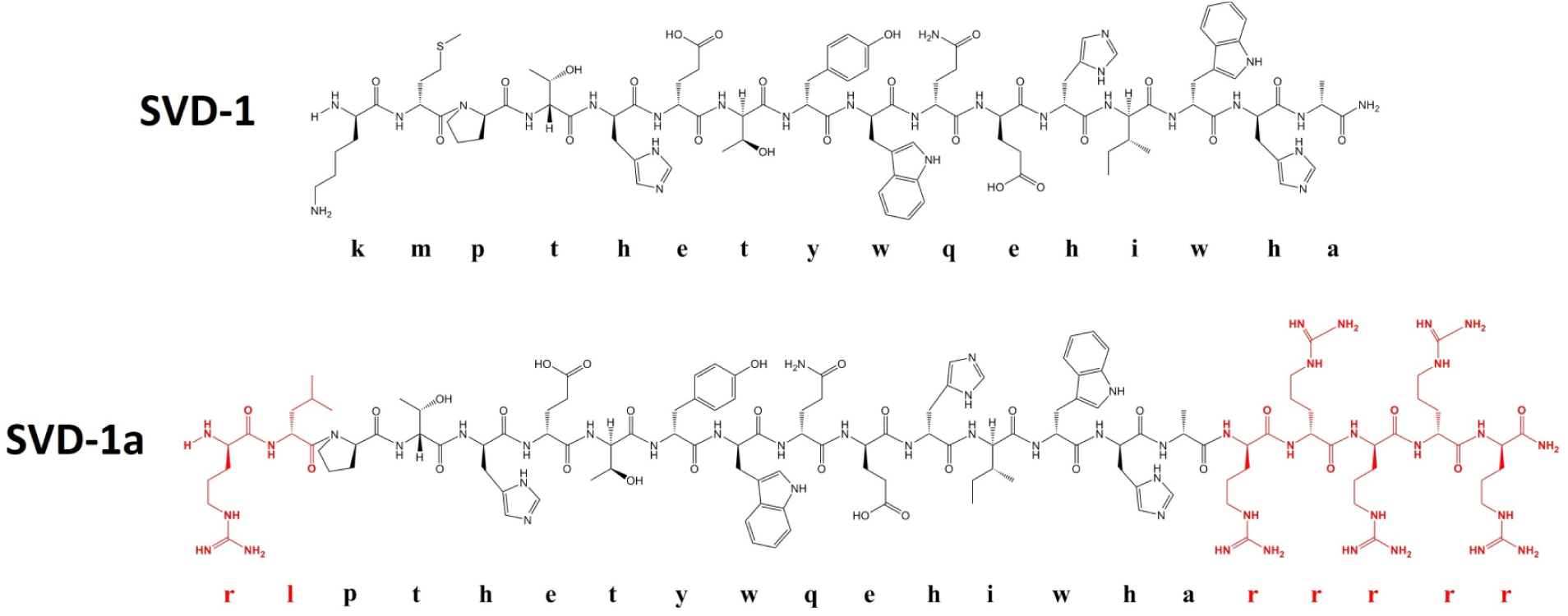
Natta projection of the D-enantiomeric lead compound SVD-1 and its first optimized derivative SVD-1a. Amino acid residues marked in red were exchanged or added for improved bioavailability, membrane penetrance and inhibitory effects.

First, we characterized binding affinities of SVD-1 and SVD-1 to α-syn monomer. SPR experiments allow the detection of kinetic values, thereby giving insights in time dependency of target recognition and complex rigidity. Prior to measurements, the SVD peptides were immobilized on a carboxyl dextran matrix surface via amino coupling and α-syn was injected as analyte in the concentration range of 30 to 500 nM (Fig. 3 A). Likewise, a control peptide that had the same amino acid composition as SVD-1 but with random amino acid sequence (SVD-1_scrambled) was immobilized. As control for SVD-1a, five arginines were added to the C-terminus of SVD-1 control peptide (SVD-1_scrambled+5r; (Fig. 3 B)).

**Figure 3:**
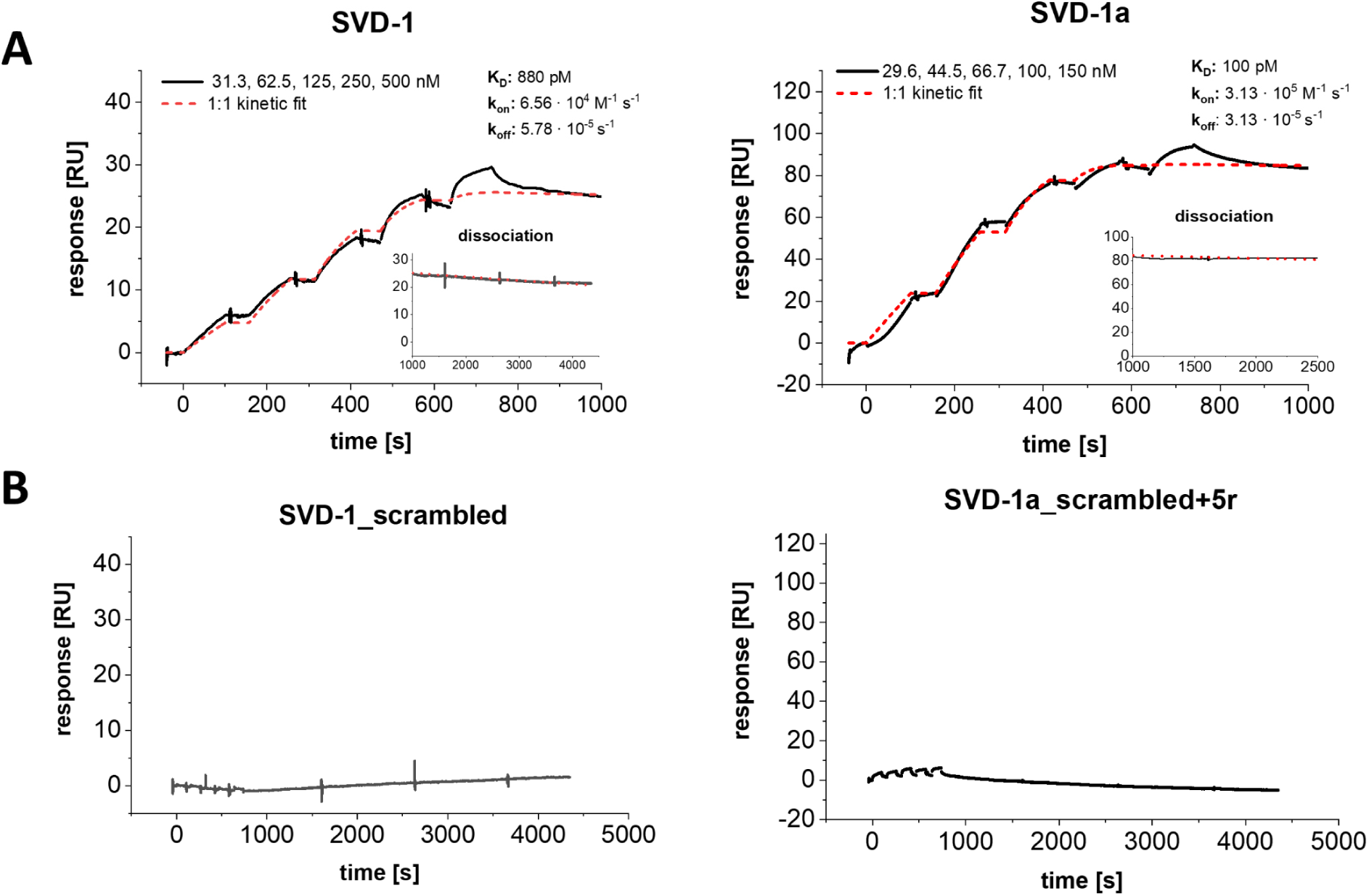
Single cycle kinetic experiment with α-syn and immobilized SVD-1 and SVD-1a and control peptides. (**A**) SVD-1 (left) and SVD-1a (right) were immobilized on a carboxyl dextran matrix via amino coupling until saturation was reached (CMD200M, Xantec, GE). a-Syn was injected for 100 s at 30 µl/min in PBS 7.4 in a serial dilution ranging from 30 to 500 nM, followed by a dissociation time of 60 min or 30 min, respectively. The experiment was performed as individual measurement. The interaction kinetics were fitted with a 1:1 kinetic interaction model: SVD-1: K_D_: 880 pM, k_on_: 6.56 · 10^4^ M^-1^ s^-1^, k_off:_ 5.78 s^-1^ · 10^-5^; SVD-1a: K_D_: 100 pM, k_on_:3.13 · 10^5^ M^-1^ s^-1^, k_off_: 3.13 · 10^-5^ s^-1^. The non-referenced signal of active and referenced surface is shown in SI Fig. 7. (**B**) SVD-1_scrambled (left) and SVD-1_scrambled+5r (right) were immobilized on a carboxyldextran matrix via amino coupling until saturation was reached (CMD200M, Xantec, GE). Full-length a-syn was injected for 100 s at 30 µl/min in PBS 7.4 in a serial dilution ranging from 15 to 250 nM, followed by a dissociation time of 60 min. Y-axis scaling was adjusted to the (A).

When applying a 1:1 Langmuir based interaction model, we obtained a K_D_ value of 880 pM with a k_on_ of 6.56 · 10^4^ M^-1^ s^-1^ and a k_off_ of 5.78 · 10^-5^ s^-1^ for SVD-1. For SVD-1a a K_D_ of 100 pM with a k_on_ 3.13 · 10^5^ M^-1^ s^-1^ and a k_off_ of 3.13 · 10^-5^ s^-1^ was identified. However, the data is not fully explained by the 1:1 model, which is evident from a deviation of the fits from the data obtained for the later injections, suggesting an additional low-affinity binding mode in the higher nM range.

The inhibitory effects on amyloid formation was further verified in ThT assays under *de novo* as well as seeded conditions with several ratios of α-syn and SVD-1or SVD-1a (SI Fig. 8). For seeding assays we incubated SVD-1 or SVD-1a together with pre-formed fibrillary (PFF) oligomers that induce fibril mass growth in the presence of monomeric α-syn. In order to obtain fibril seeds with a homogeneous size and high seeding capacity, we started from mature fibrils and treated them by harsh ultra-sonication. This yielded short fibrils with a high ratio of fibril ends per mass. Eventually remaining larger fibrils were removed by ultracentrifugation as described by Kaufmann et al. ((*32*), SI Fig. 9). Referring to their small size, we call them “PFF oligomers” throughout the manuscript. Due to their high ratio of fibril ends per mass, added monomeric α-syn is very efficiently and reproducibly elongating the seeds (*35*).

Sequence specificity of the effect was confirmed by comparing the inhibitory effects of SVD-1 and SVD-1a with the respective control peptides. The sequence-randomized control peptides had virtually no inhibitory effects, clearly suggesting that not the overall electrostatic charge or hydrophilicity were important, but the amino acid sequence of SVD-1 and SVD-1a (SI Fig. 8). In contrast to the sequence-randomized control peptide, SVD-1 confirmed its efficacy by a concentration dependent delay of aggregation onset as well as by a reduction of steady state levels under *de novo* aggregation conditions. Thus, SVD-1 inhibits primary nucleation of α-syn. Also, in contrast to the sequence-randomized control peptide, SVD-1 was able to significantly decelerate amyloid growth in the seeded environment. Thus, SVD-1 also inhibits elongation and secondary nucleation of α-syn. Similar observations were made for SVD-1a under *de novo* aggregation conditions, where at twofold molar excess aggregation was completely inhibited. When SVD-1a was present in the pre-incubation period with the PFF oligomers, a reduction of the elongation rate was observed later during the incubation period with α-syn monomers already at 5 µM SVD-1a. 20 µM or higher SVD-1a concentrations even led to the complete inhibition of the seeding capacity. 20 µM of the control peptide, SVD-1_scrambled+5r, did not yield any inhibition of seeding, again underlining the SVD-1a sequence specificity for the observed effects.

It is tempting to speculate that SVD-1a was successfully reducing the amount of PFF oligomer seeds during the 20 h pre-incubation period to explain the reduction of their seeding activity. In order to investigate this further, we incubated 100 nM monomer equivalent purified PFF oligomers with increasing concentrations of SVD-1a for 72 h and analyzed then the samples by SEC, time dependent dynamic light scattering (DLS) and atomic forced microscopy (AFM) (Fig. 4).

**Figure 4:**
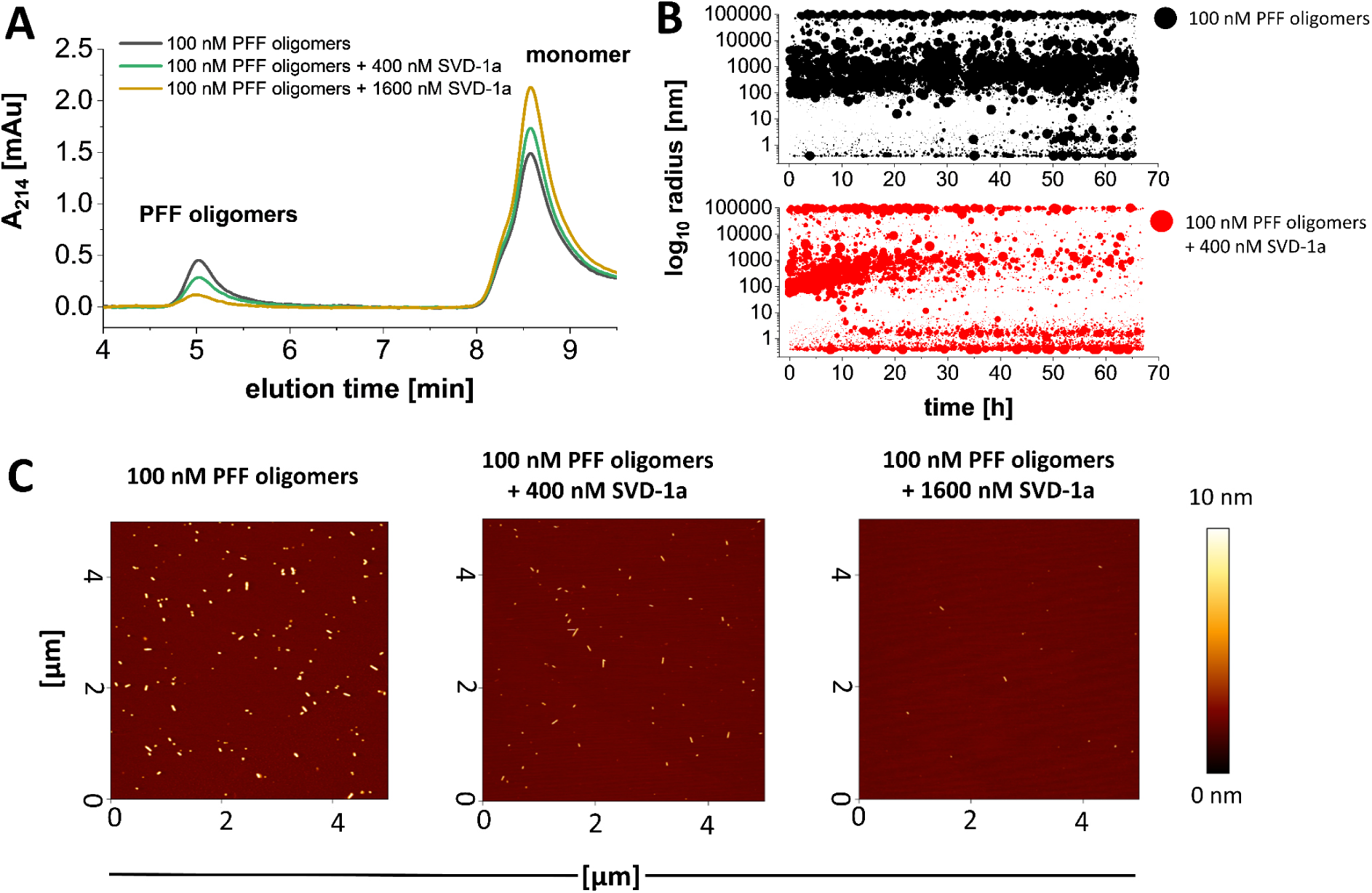
SVD-1a disassembles PFF oligomers into α-syn monomers. PFF oligomers were prepared as described previously. 100 nM monomer equivalent PFF oligomers were incubated with or without 400 nM and 1600 nM SVD-1a for 3 d at 37 °C in PBS pH 7.4. (**A**) HPLC-SEC measurement samples were injected on a Bio SEC-3 column (300 Å, Agilent, USA). The PFF oligomers elute after approx. 5 min, while α-syn monomer is detected after approx. 8.6 min. (**B**) Time dependent DLS measurements with 100 nM PFF oligomers in presence (red) and absence (black) of 400 nM SVD-1a. 1 ml sample was continuously measured in a sealed quartz cuvette at 37 °C under quiescent condition every 30 s for 72 h in a SpectroSize 300 instrument (XtalConcepts, GE). Data are shown as radius plot where the signal amplitude of each particle size is represented by the data point diameter. (**C**) For AFM analysis 5 µl of the samples as described in (A) was rescued before centrifugation and incubated and dried on a freshly cleaved mica surface followed by washing with ddH_2_O and drying using a gentle stream of N_2_. Analysis was performed using NanoWizard 3 system (J-1100, JPK BioAFM, USA), recording multiple surface sections. The sections shown in C are representative for the observed species and particle density identified on all surface sections.

The SEC analysis yielded that the PFF oligomer preparation contains a substantial fraction of monomers in equilibrium that may occur due to the proportionally high number of fibril ends as compared to mature fibrils. Incubation of the PFF oligomers with increasing concentrations of SVD-1a (here 400 and 1600 nM) resulted in an increase of monomer concentration paralleled by a decrease in oligomer concentration (Fig. 4 A). This is supported by AFM (Fig. 4 C). In addition, the time dependent DLS measurement (Fig. 4 B) shows that SVD-1a is progressively eliminating the PFF oligomers (Fig. 4 B, red, 100 to 1000 nm radius) by disassembling them into monomeric α-syn (Fig. 4 B, red, 3 nm radius). In contrast, no change of the PFF oligomer particle size radius was observed when SVD-1a is absent (Fig. 4 B, black, 100 to 1000 nm radius).

In conclusion, the analysis of the inhibitory mechanism of SVD-1a on α-syn aggregation under seeded condition has shown that (i) SVD-1a eliminates PFF oligomers in a time dependent process, (ii) stabilizes monomeric α-syn in its random coil conformation and (iii) reduces the emergence of mature fibrils.

To further verify whether SVD-1a is also able to counteract the seeding potential of soluble α-syn PFF oligomers in living cells, α-synA53T–YFP expressing cells were transfected with soluble α-syn PFF oligomers together with SVD-1a or a negative-control peptide with similar molecular weight as SVD-1a but no affinity to monomeric α-syn (Fig. 5). In the α-synA53T-YFP cell system, human α-syn with the familial A53T mutation fused to YFP is stably expressed in HEK293T cells and enables fluorescence-based detection of intracellular aggregation of endogenously expressed α-synA53T after seeding with patient-brain extracts containing PFFs as shown previously (*36–39*) or with PFF oligomers as shown here. To avoid interference with the fluorescence-based aggregate detection, we intentionally did not use fluorescence labelled SVD-1a or control peptide. Another reason was that fluorescence dyes can positively or negatively interfere with the transfection efficacy and possibly also with the subcellular localization of the respective peptide.

**Figure 5:**
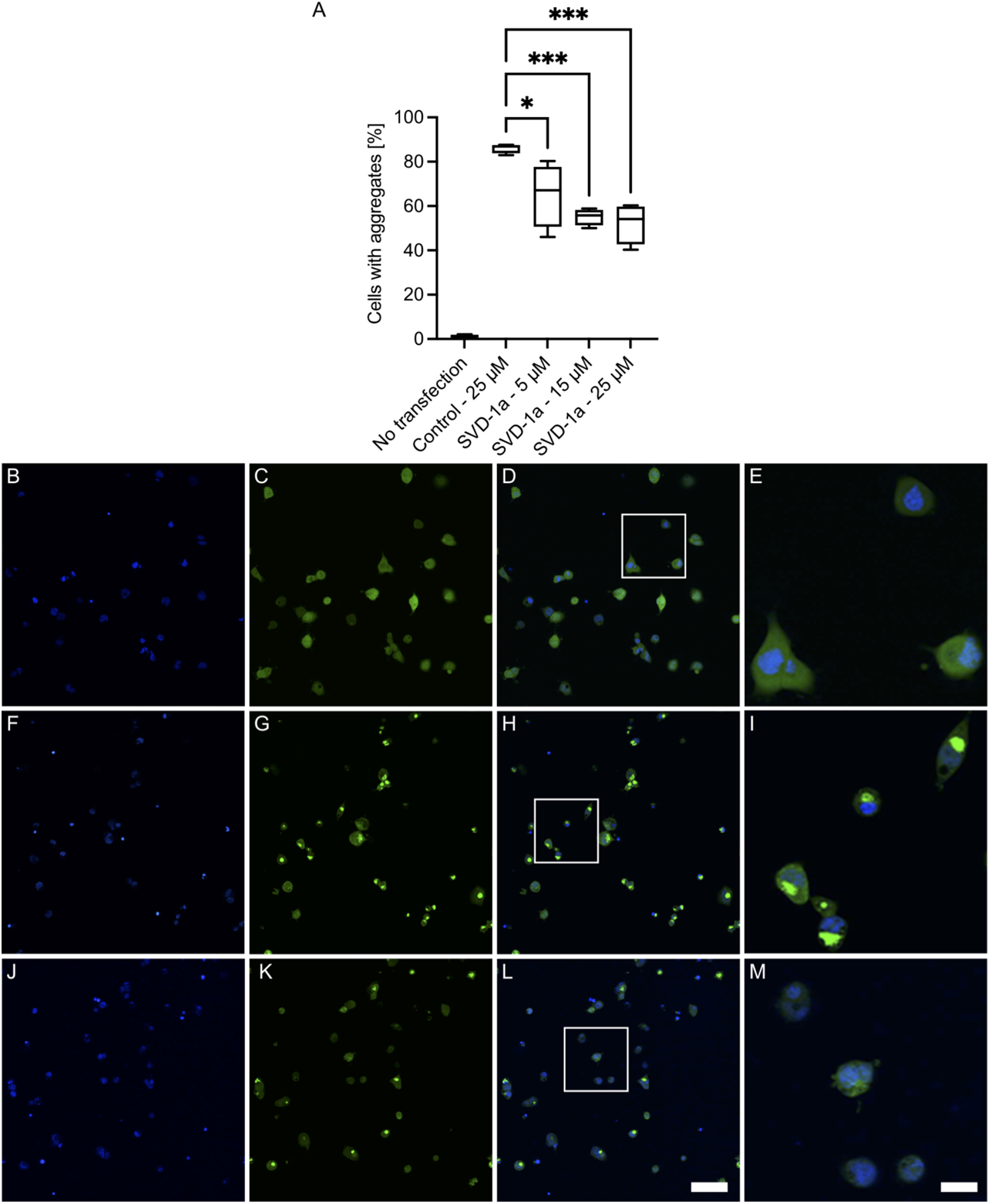
SVD-1a inhibits seeded α-syn aggregation in cells. To validate the inhibitory effect of SVD-1a in cells, we used the α-synA53T–YFP cell system, which stably expresses human α-syn with the familial A53T mutation fused to YFP in HEK293T cells and enables fluorescence-based detection of aggregates that show up as highly fluorescent spots within the cells clear above background, intracellularly after seeding with PFF oligomers. In order to avoid interference of the peptide with the uptake of PFF oligomers, we performed a two-step transfection starting with a first transfection of peptide and a second transfection with PFF-oligomers. SVD-1a but not the negative control peptide (kgvgnleyqlwalegk-NH2) inhibited α-syn aggregation in αSynA53T–YFP cells in a dose-dependent manner (**A**). To quantify the number of cells with aggregates in our images and to avoid experimental bias, we used a fully automated algorithm for image analysis. Significance was calculated using one-way ANOVA followed by Dunnett’s multiple comparisons test. One asterisk denotes p values less than 0.05 and three asterisks a p value less than 0.001. Error bars indicate standard deviation. In contrast to unseeded cells, which did not harbor any α-syn aggregates at day three after plating (**B–E**), seeding with soluble α-syn PFF oligomer induced aggregation in 85 % of cells treated with a negative-control peptide (25 µM) that does not bind to α-syn (**F–I**). Treatment with increasing concentrations of SVD-1a, here imaged at 25 µM, led to a concentration-dependent reduction in the number of cells with aggregates (**J–M**). Panels B, E, and H show nuclei stained with Hoechst 33342. Panels C, F, and I show αSynA53T–YFP fluorescence. Panels D, G, and J show merged images. The scale bar in L represents 100 µm and applies to panels B-D, F-H, and J-L. Panels E, I, and M represent magnifications of the insets in D, H, and L, respectively the scale bar in M applies to panels E, I and M and represents 25 µm. The viability of the cells was not reduced by transfection with seeds or in SVD-1a as shown in SI Fig. 12. Confirmation that highly fluorescent aggregates contain α-syn is shown in SI Fig. 10 by co-immunostaining with syn-211 antibody, anti-α-syn (phospho S129) antibody and 5G4 antibody.

Seeding with α-syn PFF oligomers in the presence of a negative-control peptide induced aggregation in 85 % of the cells (Fig. 5 A, E–G). We verified that the induced intracellular, yellow-fluorescent aggregates consist of α-syn by immunofluorescence staining with three different antibodies recognizing aggregated α-syn, α-syn phosphorylated at serine 129, and total α-syn (SI Fig. 11). When SVD-1a was transfected into the cells, a concentration-dependent reduction of aggregate-positive cells was observed resulting in 64 % and 57 % aggregate-positive cells using 5 and 25 µM of SVD-1a, respectively (Fig 13 A, H–J). Also, the viability of PFF oligomer-seeded α-synA53T–YFP cells was not significantly reduced in the presence of any of the two peptides (SI Fig. 12. SVD-1aCys_Alexa647 is crossing the membrane without transfection (SI Fig. 13). These results demonstrate that SVD-1a inhibits PFF oligomer-induced aggregation of α-syn in the intracellular environment without inducing cytotoxicity. Moreover, in PC12 cells a reduction of PFF oligomer cytotoxicity was verified in presence of SVD peptides using cell viability assay (SI Fig. 14).

SVD-1 and SVD-1a have been selected and developed to specifically bind α-syn monomers in order to stabilize α-syn in its monomeric conformation, which is well known to be a classic example of an intrinsically disordered protein (IDP). The constant and fast sampling of a large conformational space gives IDPs the structural plasticity and adaptability to interact with and control multiple binding partners at the same time. IDPs have very characteristic NMR spectra. The amide protons of the protein backbone are solvent exposed and not involved in typical secondary structural elements like β-sheets or α-helices. This and the high mobility of the protein backbone and the side chains on a very rapid time scale limit the chemical shift dispersion of IDPs more or less to the time averaged random coil chemical shifts of the respective amino acid residues of the protein in aqueous solution. This is the reason, why the amide protons of IDPs have a typical chemical shift dispersion of only 0.7 ppm. In contrast, the amide protons of globularly folded proteins have a chemical shift dispersion of up to 4 ppm, and the individual chemical shift of the amide proton is dependent of its involvement in a hydrogen bond and whether its chemical environment contributes more shielding or de-shielding contributions, both of which are not averaged out over time as in IDPs (*40, 41*).

Of course, we were very curious to investigate, whether SVD-1 and SVD-1a have a significant impact on the IDP conformation of α-syn monomers, given their high affinity based on SPR. The highly dynamic and transient interactions may for each amide proton and ^15^N-nucleus lead to many different shielding and de-shielding events in their chemical environment that become zero, when averaged over the NMR time scale. The probability, however, that each chemical shift change is averaged to exactly zero, is very low. Thus, we investigated the chemical shift changes upon binding of SVD-1a to α-syn monomers at the highest available field. Figure 6 A shows the superposition of the ^1^H-^15^N HSQC NMR spectra of 25 µM ^15^N-labeled full-length α-syn in absence and presence of 25 µM SVD-1a. Careful and automated peak analysis revealed that there are indeed small chemical shift changes that are shown α-syn sequence specifically in Figure 6 C with some examples displayed in Figure 6 B. Overall, the chemical shift changes are very small, with most residues showing significant chemical shift changes located in the C-terminal region but also residues in other parts of α-syn are affected.

**Figure 6:**
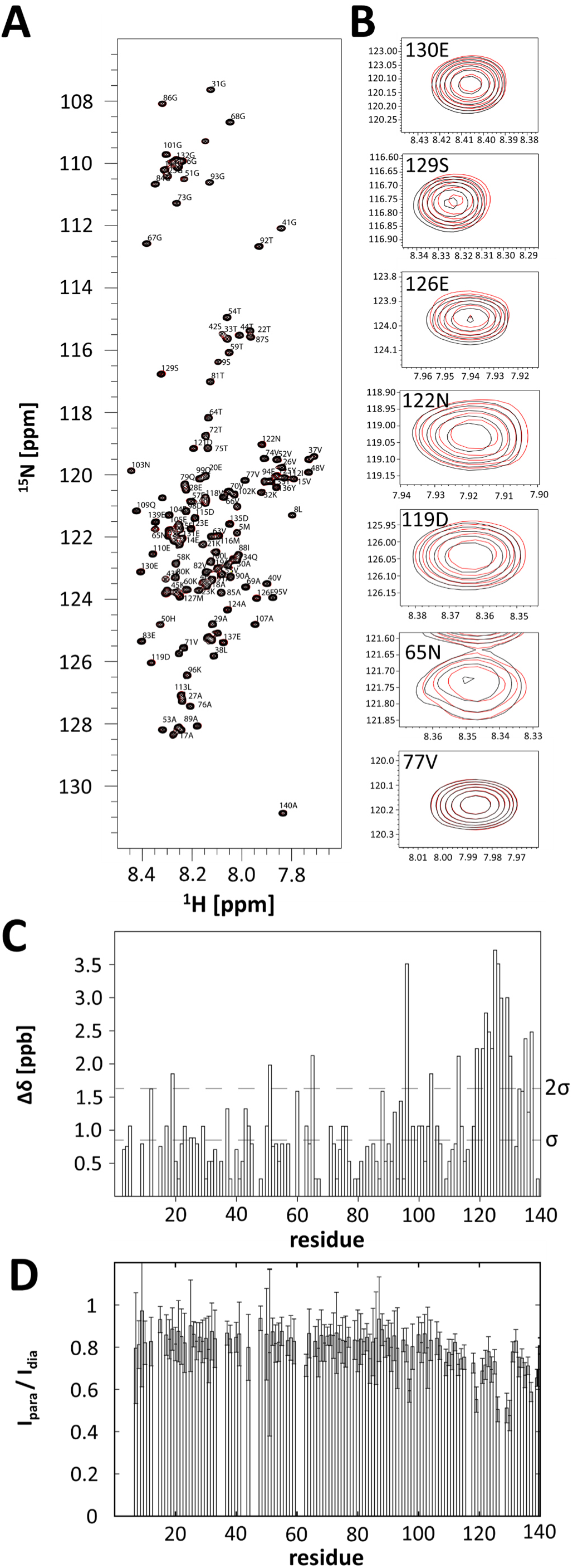
NMR analysis of ^15^N-labeled α-syn interacting with SVD-1a (not isotope labeled). **(A)** Overlay of two-dimensional ^1^H-^15^N HSQC spectra of 25 µM ^15^N α-syn in the presence (red) and absence (black) of an equimolar amount of SVD-1a. **(B)** Enlargement of several resonances in the spectra shown under (A) that show small chemical changes for residues in the presence (red) relative to the absence (black) of SVD-1a (130E, 129S 126E, 122N, 119D, 65N); for comparison, resonance 77V is shown that does not show any chemical shift change. **(C)** Residue-specific absolute NMR chemical shift changes in the spectra of ^15^N α-syn in presence of SVD-1a relative to the absence of SVD-1a. The standard deviation, σ, as well as the 2-fold standard deviation, 2σ, of the distribution of observed chemical shift changes are indicated as dashed lines. **(D)** NMR PRE intensity ratios of ^15^N α-syn in the presence of the paramagnetically-labeled SVD-1a. Residue-specific intensity ratios, I_para_ / I_dia_, of the cross-peak intensities in the two-dimensional ^1^H-^15^N NMR spectra of the paramagnetic vs diamagnetic sample are shown. The lower the intensity ratios, I_para_ / I_dia_, the closer the proximity of the paramagnetically-labeled SVD-1a to the respective residue of α-syn. An intensity of one would indicate the absence of interactions. Data point to a bit more pronounced (transient) binding interaction of SVD-1a with residues in the C-terminal region of α-syn, as compared to the average effect on residues in the remaining N-terminal region.

To obtain more information, which parts of the α-syn molecule are involved in the highly dynamic and transient interaction, we applied paramagnetic relaxation enhancement (PRE) NMR experiments. Such NMR-PRE data have proven insightful for the study of binding interactions of amyloid-β recently (*42*).

Residue-specific NMR PREs intensity ratios, I_para_ / I_dia_, were recorded for ^15^N-labeled (NMR-visible) α-syn in the presence of SVD-1a, with a paramagnetic spin-label covalently attached at its C-terminus (Fig. 6 D). Decrease of intensity ratios point to proximity of the paramagnetic spin-label of SVD-1a to the respective residue of α-syn. Indeed, we observed an overall decrease of intensity ratios, I_para_ / I_dia_, in the spectra (paramagnetic vs diamagnetic sample) for practically all residues, in the order of 15 to 20 % and with an increased reduction for the C-terminal residues of α-syn (Fig 6 D). Strikingly, this coincides with the residues showing the “largest” of the very small chemical shift changes (Fig. 6 B) that were most prominent in the C-terminal region. The apparently weak transient interactions with the N-terminal 108 α-syn residues, as observed by the NMR-PRE data, are in their accumulated sum effect relevant and impactful. This is indicated by the observation that the *de novo* aggregation not only of full length α-syn is efficiently inhibited by SVD-1 and SVD-1a, but also that of the C-terminal deletion mutant of α-syn, α-syn (1–108) (SI Fig. 15).

The PRE data together with the absence of large chemical shift changes are in-line with lowly populated transient binding interactions, presumably due to an on/off hopping of SVD-1a to α-syn occurring on a very fast time scale. Hence, the binding mode is potentially best described by an IDP – IDP interaction with multiple dynamic binding sites, whereas both partners retain their disordered structure (*12, 43*). Such fuzzy complexes have been described previously (*44–50*). In some cases no chemical shift changes have been reported at all (*44*). For several of those cases, NMR paramagnetic relaxation enhancement (PRE) measurements revealed transient interactions between the protein and the binding partner (*44–50*).

To investigate in further detail how this interaction mode functions under aggregation promoting conditions, we performed *de novo* aggregation in presence and absence of the peptide and analyzed the endpoint samples by circular dichroism spectroscopy (CD) and AFM.

CD secondary structure analysis (Fig. 7 B) of replicate samples without ThT shows that incubation of monomeric α-syn results in a shift from random coil (Fig. 7 B, black line) to beta-sheet (Fig. 7 B, blue line) CD spectrum, typical for α-syn fibrils (*51*). However, in the presence of SVD-1a, while the overall signal is slightly reduced, no spectrum shift towards a beta-sheet spectrum was observed (Fig. 7 B, grey line). In combination with the ThT measurements, the CD measurements clearly indicate that the majority of monomeric α-syn that is present at the start point of incubation, is retained in its monomeric random coil form when SVD-1a is present and is only aggregating into β-sheet rich fibrils when SVD-1a is not present. These observations are confirmed by AFM pictures of the samples taken from the incubation endpoints in the ThT assay of figure 11 A (Fig. 7 C). Here, the sample without SVD-1a shows mature fibrils (Fig. 7 C, left). The AFM pictures from samples that contained SVD-1a reveal small particles that are artifacts due to the drying of the samples (Fig. 7 C, right) before imaging. However, they clearly do not contain any fibrils, supporting that SVD-1a was able to inhibit fibril formation as already demonstrated by the ThT experiments.

**Figure 7:**
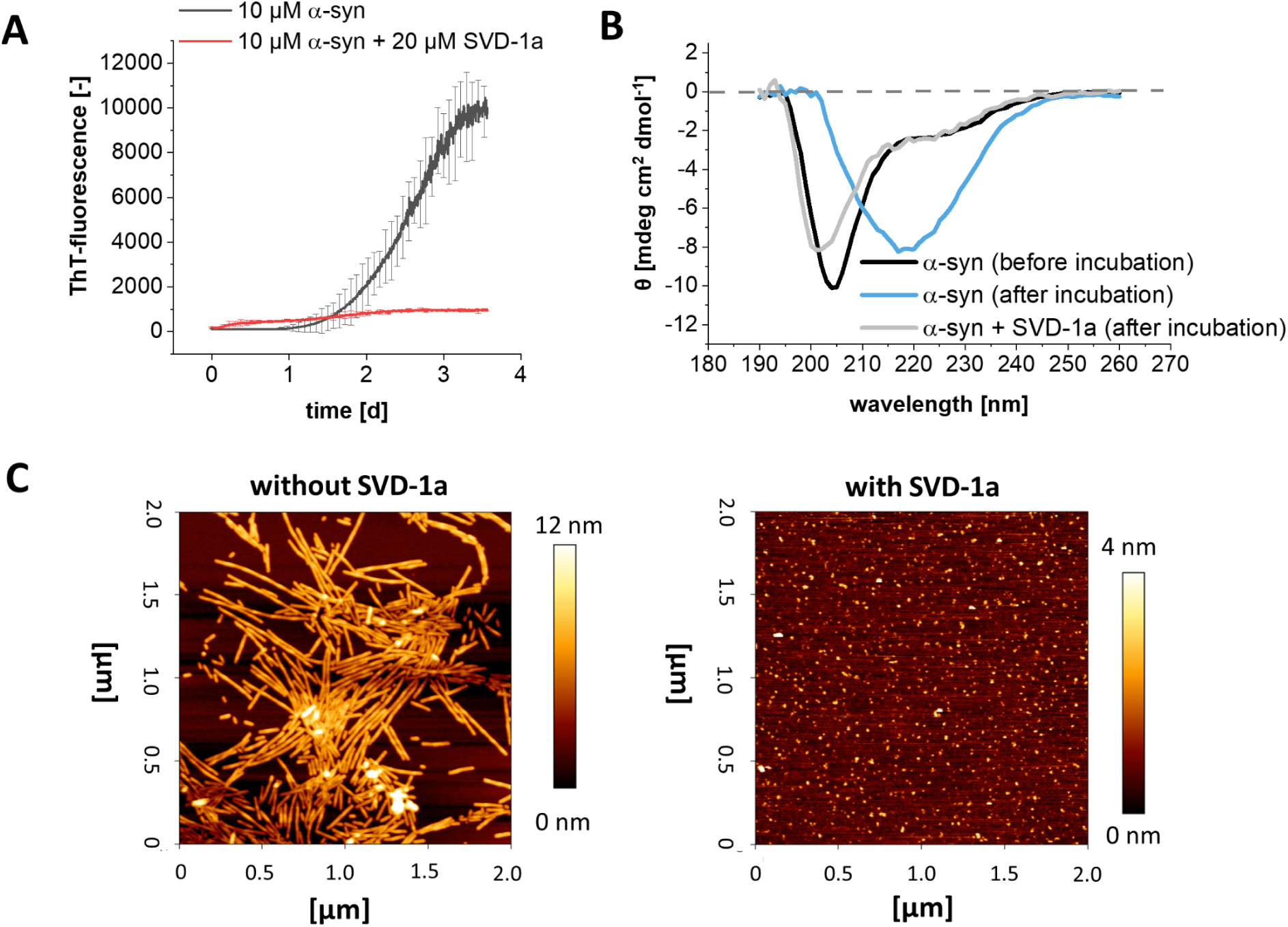
*De novo* aggregation analysis of α-syn in the presence and absence of SVD-1a using ThT, DGC, CD and AFM. **(A)** *De novo* ThT assay of 10 µM α-syn with and without 20 µM SVD-1a. ThT fluorescence progression was measured in a 96-well non-binding half-area plate (Corning, USA) with a Fluorostar platereader (BMG labtech, GE) at λex = 448 nm and λem = 482 nm with 300 rpm continuous orbital shaking between reads. Data are shown as mean values with ± SD (n = 5). (**B**) CD secondary structure analysis of *de novo* aggregation samples. Samples were incubated as described in (A) without added ThT (n = 3) and subsequently pooled for CD analysis. Far-UV ellipticity of the samples was measured in a quartz cuvette (l = 10 mm) in a J-1100 CD-spectrometer (Jasco, GE). In addition to (A), a sample with 20 µM SVD-1a alone was incubated under identical conditions and later used as reference for the sample with α-syn and SVD-1a. For this sample (α-syn + SVD-1a (after incubation)) the SVD-1a reference subtracted CD spectrum is shown. (**C**) Samples from (A) were isolated directly after incubation and diluted in PBS pH 7.4 to a final concentration of 1 µM α-syn monomer equivalent. 5 µl diluted sample was incubated and dried on a freshly cleaved mica surface followed by washing with ddH_2_O and drying using a gentle stream of N_2_. Analysis was performed using NanoWizard 3 system (J-1100, JPK BioAFM, USA), recording multiple surface sections. The section shown in (C) are representative for the observed species identified on all surface sections.

In order to integrate the results shown into a mechanistic model (Fig. 1) of a therapeutic mode of action, the following considerations are crucial: only very small chemical shift changes were identified for unbound vs bound α-syn, indicating that the presence of the compounds did not significantly change the overall α-syn conformation. At first glance, this may seem counter-intuitive when compared to the classical receptor-ligand interaction studies, where one would expect chemical shift changes for the residues close to the “binding pocket”. However, the binding mode of IDPs, such as α-syn, may substantially deviate from such classical picture (*52, 53*). Thus, we carried out NMR experiments with ^15^N isotope-labelled α-syn and SVD-1a with a paramagnetic label attached. When applying paramagnetic relaxation enhancements (PRE) experiments the PRE-label associated with the peptide decreases the peak intensities around the binding site. As our PRE measurements indicate (Fig. 6 D), SVD-1a interacts with the entire α-syn monomer, with the C-terminal part showing the strongest interaction. PRE data indicate a transient and highly dynamic interaction.

The binding data would be consistent with the following scenario: SVD-1 and SVD-1a encounter α-syn in various conformations. Each conformation of this conformational ensemble is transient and sparsely populated, with conformations interconverting among each other on fast time scales. In sum, the accumulated high number of sparsely populated binding events will result in the observed macroscopic high affinity binding constant. Note, that the observed overall binding will not necessarily induce large chemical shift changes for the individual residues (*44, 45*), but leads to the observed NMR PRE effects. This is a binding mode, which comes closest to the envisioned mode of action for SVD-1 and SVD-1a, namely to stabilize α-syn monomers in their highly dynamic and flexible IDP conformation.

For the majority of the α-syn anti-aggregation compounds that have been developed over the last years, an essential dogma was the avoidance of a direct interference with the physiological monomeric form of α-syn in order to exclude effects that might influence the physiological function of the target protein (*54, 55*). The following data support the conclusion that SVD-1a binds to the α-syn monomer, stabilizes it in its IDP-like conformation, and keeps it monomeric in its physiological IDP conformation. Incubation of 25 µM α-syn with 25 µM SVD-1a for 4.8 h at 10 °C for the NMR experiments shown in Figure 6 A did not yield any signs of signal loss due to precipitation. Similarly, for the PRE NMR experiments (Fig. 6 D), the observed amide cross peak intensity reduction due to the PRE label interaction with α-syn residues, was fully rescued upon addition of ascorbate to quench the PRE label (which was the diamagnetic reference experiment). Thus, SVD-1a is not sequestering α-syn monomers into any other conformation or state. SVD-1a is rather stabilizing α-syn monomers in their IDP conformation by the free binding energy underlying the high affinity demonstrated by the SPR measurements (Fig. 3). The strong binding of SVD-1a to α-syn monomers is not influencing its IDP conformation, suggesting that its physiological role in the cell might not be affected or limited by SVD-1a. Figure 1 A illustrates, why the stabilization of α-syn monomers by the free binding energy of SVD-1a, is also disassembling already existing PFF oligomers (Fig. 1 B), just because SVD-1a bound α-syn monomers are thermodynamically more stable than the α-syn building block conformation in oligomers. This strongly supports the proposed mode of action of SVD-1a. Disassembly of PFF oligomers by SVD-1a as demonstrated by SEC analysis of PFF oligomers incubated with SVD-1a by the observed increase of α-syn monomer concentration paralleled by the decrease of the PFF oligomer concentration, in an SVD-1a concentration dependent manner, also verified by DLS (Fig. 4). All these results support the mechanistic model for SVD-1a’s mode of action as described in Fig. 1.

Taken together, the oligomer-elimination assay as well as the assays for which PFF oligomers were used as the pre-formed aggregate species (cell viability assay, intracellular aggregation assay, seeded aggregation assay) show that the compounds are able to eliminate soluble α-syn aggregates independent of their overall structural assembly. This result appears to be in agreement with the intended anti-prionic mode of action, where the compounds stabilize the physiological IDP-like monomer structure, thereby destabilizing and disassembling the toxic aggregates. This allows the mode of action to be independent of the specific conformation of specific toxic aggregate assemblies. The physiological solution structure of monomeric α-syn remains the same *in vitro* and *in vivo*, irrespective of the localization (*56*). This makes monomeric α-syn a more attractive target, since this mode of action is independent of the final form of the toxic component, and thus independent of any prion strain.

These results are a promising starting point for further development of an all-D-enantiomeric peptide compound is even disassembling already existing aggregates. Since the compounds presented here are predominately interacting with the physiological active monomeric form of α-syn, future studies will also address the preservation of its physiological functionality in presence of the compounds. In addition, future efforts will deal with the investigation of the compound’s blood-brain-barrier penetrance and pharmacokinetic profile to show the transferability of the anti-prionic mode of action *in vivo*.

## CONFLICT OF INTEREST

MS, JM and DW are co-inventor of patents covering the composition of matter of SVD-peptides. DW and AW are co-founders and shareholders of the company “Priavoid GmbH, Düsseldorf, Germany”, which is planning to further develop the SVD-peptides. They declare that this has not influenced to evaluation and interpretation of experiments. All other authors declare no competing interest.

## AUTHOR CONTRIBUTIONS

**Conceived and designed the project outline**: MS, WH, AW, GT, JK, JM and DW. **Planned and performed experiments**: Phage display selection (MS, WH, JM, DW); ThT screening assays (MS); *de novo* and seeded ThT assays (MS); peptide optimization (MS, JM, DW); surface plasmon resonance (MS); circular dichroism spectroscopy (MS); dynamic light scattering (MS); atomic forced microscopy: (MS, TK), nuclear magnetic resonance spectroscopy (MS, MS^2^, NAL, LG), intracellular seeding assay (MS, MV, SAS, GT); PFF oligomer elimination (MS); cell viability assay (MS, MT, JK); **wrote the manuscript:** MS, NAL, GT, JM and DW; **contribution for revision**: LNS, TP. All authors contributed to the manuscript.

## DATA AVAILABILITY STATEMENT

The datasets generated during and/or analysed during the current study are available from the corresponding author on reasonable request.

## RESEARCH FUNDING

The project received funding by the Michael J. Fox Research Foundation (Grant-ID: MJFF-000934).

## ACKNOWLEDGEMENTS

The authors acknowledge access to the Jülich-Düsseldorf Biomolecular NMR Center that is jointly run by the Forschungszentrum Jülich and Heinrich-Heine-University Düsseldorf. We acknowledge Abishek Cukkemane for help with time dependent DLS experiments as well as Lena Mangels for help with AFM measurements.

---

**SI Figure 1:**
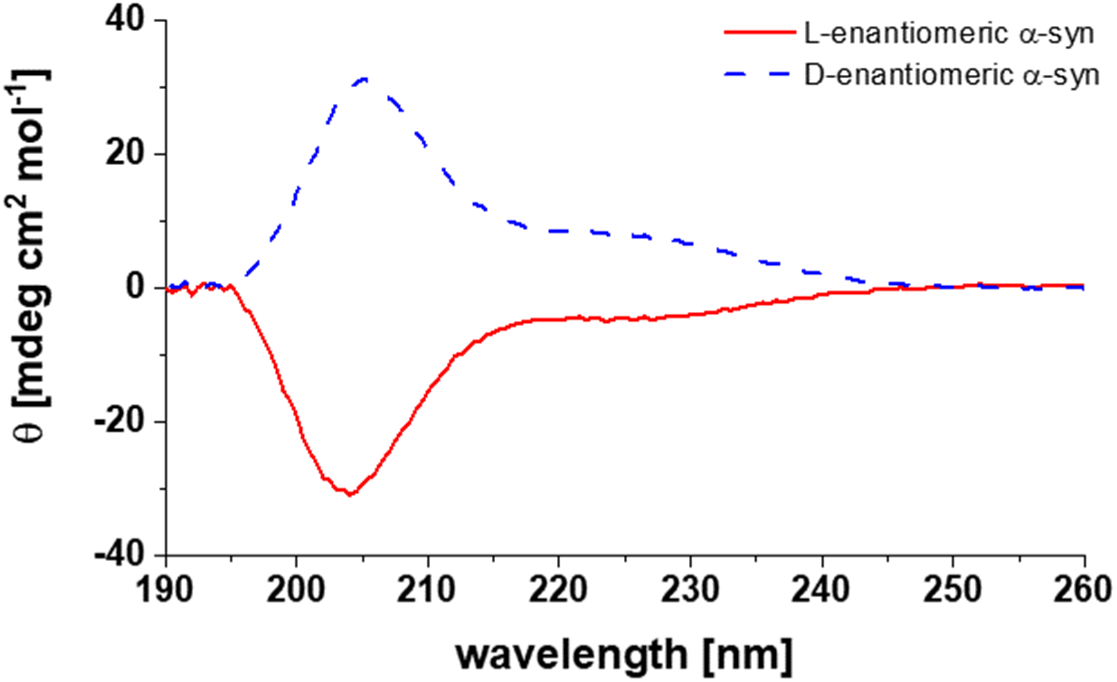
CD-measurement of recombinant L- and synthetic D-enantiomeric full-length α-syn. The D-enantiomeric C-terminally biotinylated synthetic α-syn was used as selection target. Measurements of 20 µg ml^-1^ protein in 50 mM NaPi pH 7.4 were performed as described in the method section.

**SI Figure 2:**
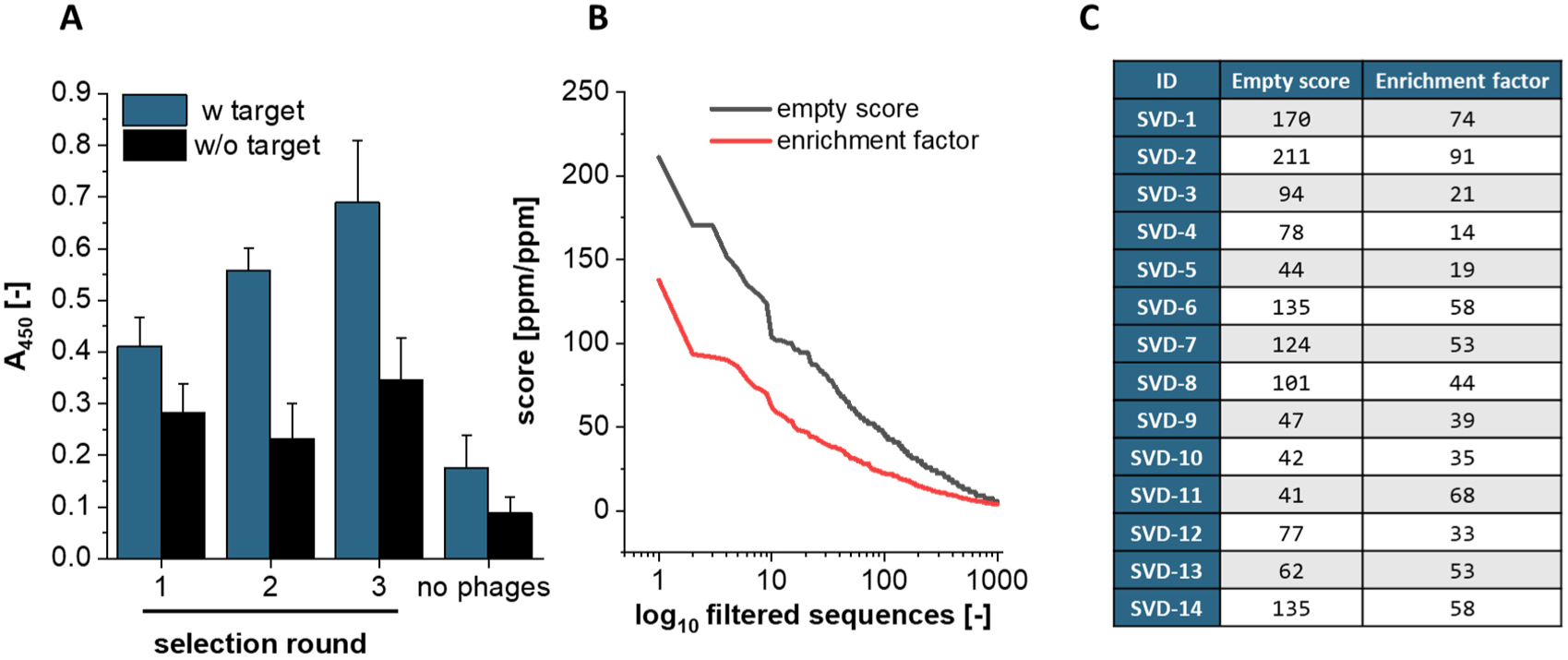
Results of the mirror image phage display selection and NGS analysis. Mirror-image phage display and NGS analysis was performed as described as follows: The full-length D-enantiomeric α-syn target protein (SI Fig. 1) was immobilized by directed coupling on a streptavidin derivatized surface via a C-terminal biotin to ensure a homogenous interaction surface during biopanning with the 16-mer M13 phage library. In addition to incubation of the phage library with the target protein (TS = target selection), an identical parallel selection, which lacks the target immobilization (ES = empty selection), was performed. Each purified phage suspension resulting from one TS round was additionally incubated on a surface without target (DC = direct control). Enrichment of target binding phages was verified after three selection rounds by direct detection of the phage coating protein in an ELISA set-up. The purified DNA of the phage suspensions was analyzed by NGS resulting in 0.9 – 1.1 Mio total sequences per sample. Target related sequences were identified after normalization by filtering specifically for those sequences whose number increased during selection rounds (TS3 > TS2 > TS1 > library) and occurred with a higher frequency in TS than in the corresponding controls that miss the target (TS1 > ES1, TS2 >ES2, TS3 > ES3, TS2 > DC2, TS3 > DC3). Thus, the sequence pool was narrowed down to 140,000 sequence variants with ∼ 4,000 showing more than one appearance. The sequences were ranked by a scoring system based on enrichment (enrichment factor = TS3/ library) and target related occurrence (empty score = TS3/ ES3), which allowed the identification of the sequence variants with the highest probability for a target interaction. In addition, clustering of the filtered sequences allowed the identification of motifs and possible cluster extensions. Based on these results, 14 D-enantiomeric synthetic peptides were further analyzed in in vitro experiments. (**A**) Enrichment ELISA of phage suspensions after amplification. 2.5 · 10^11^ phages were incubated with or without 20 pmol/well immobilized D-enantiomeric full-length α-syn target protein (n = 3). (**B**) Scoring values for enrichment (empty score = TS3/ ES3) and target specificity (enrichment factor = TS3/ library) of the filtered sequences resulting from NGS sample analysis. (**C**) List of sequences that were selected for synthesis based on NGS-scoring, empty score and enrichment factor. These sequences were screened for aggregation delaying effects in ThT-assay as shown in SI Fig. 4.

**SI Figure 3:**
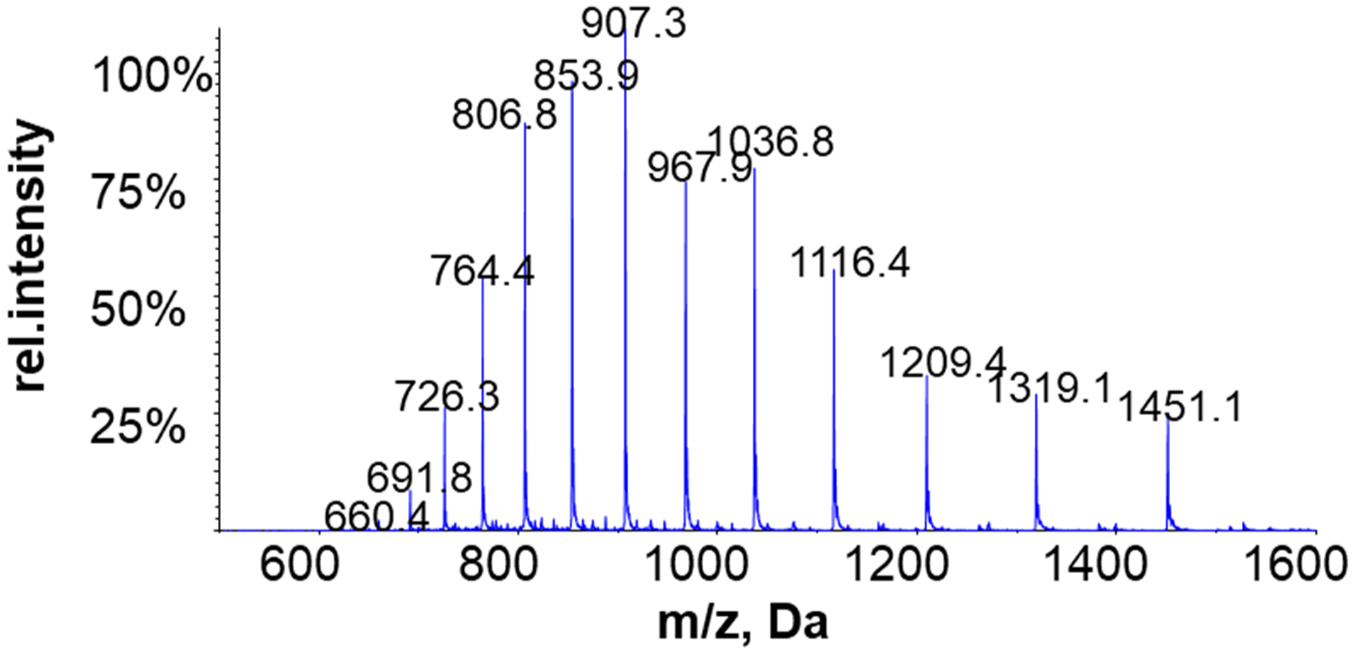
Mass spectrometry spectrum of recombinant α-syn. An HPLC-MS method with ESI as an ionization mode with a Triple Quadrupole Qtrap6500 instrument (ABSciex, GE) coupled with an Agilent 1260 HPLC system (Agilent, GE) was used. The reversed-phase column Thermo, Accucore-150-C4 (100*4.6 mm 2.6 µm) was used with the following gradient: the gradient started with 5 % B for 5 min, followed by a 15 min ramp from 5 % to 9 % B followed by a hold of 95 % B for 5 min and a re-equilibration of 10 min with 5 % B. [(A: 0.1% v/v formic acid in H2O), (B: 0.1 % v/v formic acid in ACN)] at 30°C and an injection volume of 10 µl. Mass deconvolution was performed with ESIProt (*57*).

**SI Figure 4:**
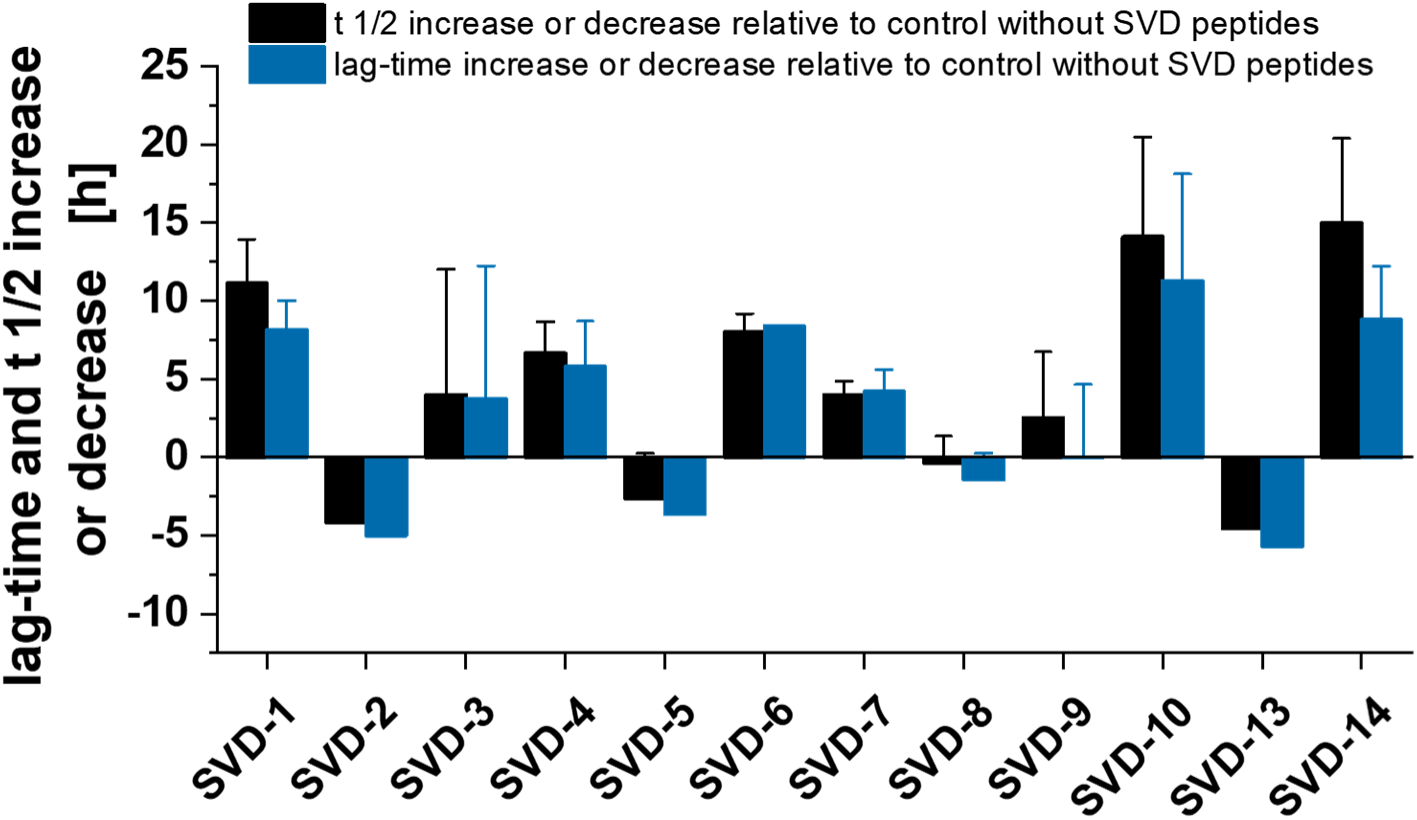
Thioflavin-T assay with α-syn and D-enantiomeric 16-mer peptides SVD-1 to SVD-14. Monomeric α-syn (SI Fig. 3) was incubated with or without a 3-fold molar excess of the D-peptides. Peptides SVD-11 and -12 were found to be insoluble in aqueous buffer and organic solvents and were therefore excluded from further analysis. The ThT positive aggregation was fitted with a symmetric Boltzmann fit and t ½- and lag-time were calculated accordingly. Since ThT signal is not always correlating with absolute amyloid mass (since different inhibitors can induce the formation of different fibril polymorphs) we decided not to evaluate the steady state in the early screening phase but rather concentrate on peptide induced t ½ - and lag-time shifts. 50 µM recombinant α-syn was incubated with or without a three-fold molar excess of the respective D-peptide in PBS pH 7.4 at 37°C. ThT fluorescence progression was measured in a 96-well non-binding half-area with a Fluorostar plate reader at λex = 448 nm and λem = 482 nm. Samples were shaken at 300 rpm for 30 s per cycle using orbital shaking mode t ½- and lag-time increase or decrease were calculated after Boltzmann sigmoidal fitting. The t ½ is given by the inflection point of the fit and the lag-time was approximated using the formula [lag-time = t ½ - 2dx] where dx is defined as the slope of the fit at x = t ½. t ½ and lag-time increase or decrease were determined based on the differences between samples with and without SVD peptide, whereas the 0-value represents the reference control without inhibitor. Mean data are shown with ±SD (n = 3). For α-syn without peptide the following t ½- and lag-times were determined: t ½ 50.5 h (PBS) and 50.1 h (PBS + 2.5 % DMSO); lag-time: 38.2 h (PBS) and 38.9 h (PBS + 2.5 % DMSO). Non-normalized aggregation curves are provided in SI Fig. 5. Seven out of twelve D-peptides delayed α-syn aggregation (SVD-1, 3, 4, 6, 7, 10, 14), whereas three D-peptides promoted the aggregation onset (SVD 2, -5, -13) and two D-peptides did not show significant impact on the aggregation (SVD 8 and -9). Among the peptides that delayed the aggregation, SVD-1, -4, -6, -10 and -14 showed the strongest inhibitory effects ranging from + 5 to 15 h t ½ - and lag-time shifts. Next, to verify the concentration dependency of the inhibitory effect, α-syn was incubated with increasing peptide concentrations, starting with equimolar concentration and continuing with a three-fold and five-fold molar excess, respectively (SI Fig. 6).

**SI Figure 5:**
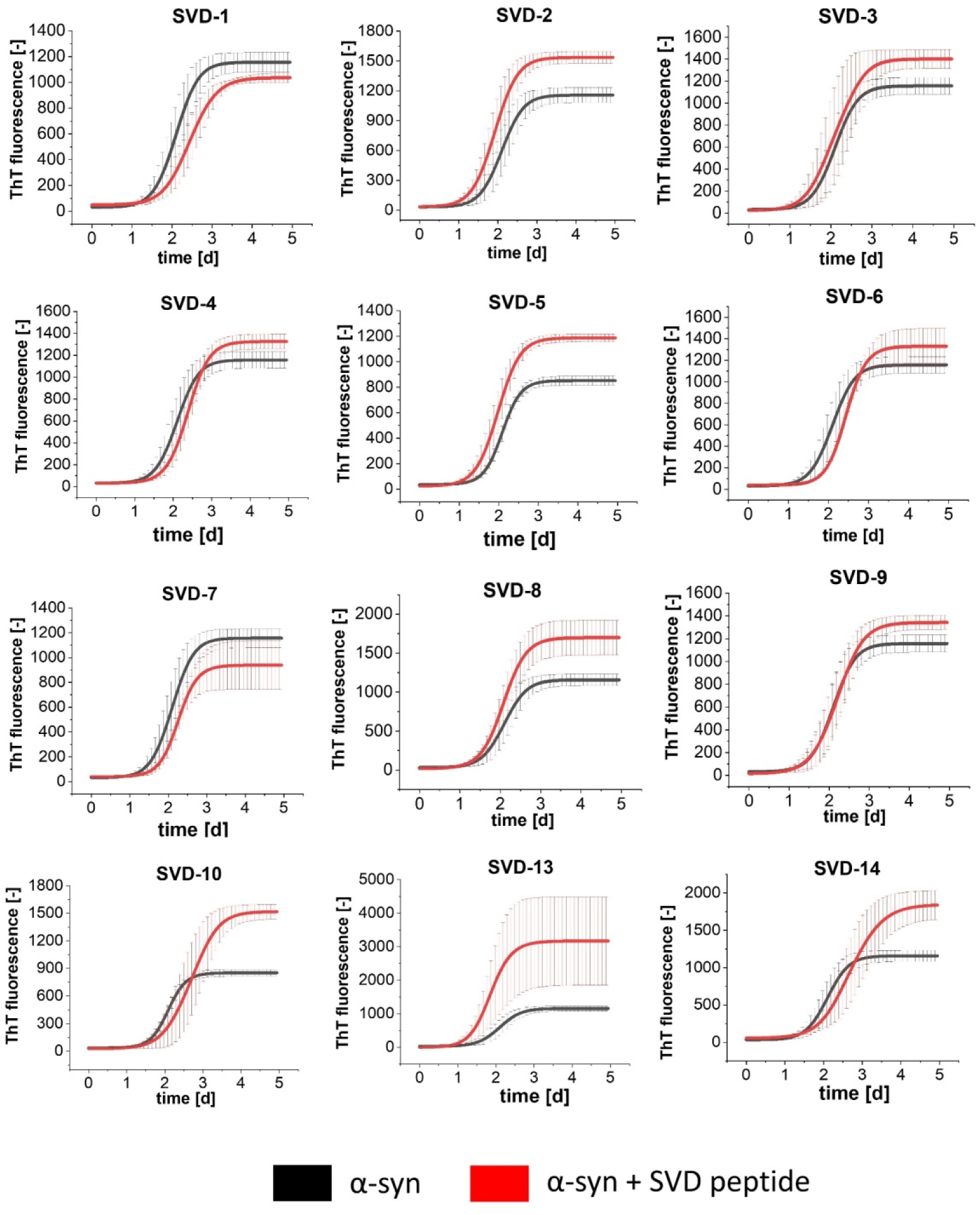
Non-normalized ThT data of *de novo* aggregation assay screen with α-syn and SVD peptides 1 to 14 as shown in SI Fig. 4. 50 µM recombinant full-length L-α-syn was incubated with or without a three-fold molar excess of the respective D-peptide in PBS pH 7.4 at 37°C. ThT fluorescence progression was measured in a 96-well non-binding half-area with a Fluorostar plate reader at at λex = 448 nm and λem = 482 nm. Samples were shaken at 300 rpm for 30 s per cycle using orbital shaking mode t ½- and lag-time shifts were calculated after Boltzmann sigmoidal fitting. Mean data are shown with ±SD (n = 3). For α-syn without peptide the following t ½- and lag-times were determined: t ½ 50.5 h (PBS) and 50.1 h (PBS + 2.5 % DMSO); lag-time: 38.2 h (PBS) and 38.9 h (PBS + 2.5 % DMSO).

**SI Figure 6:**
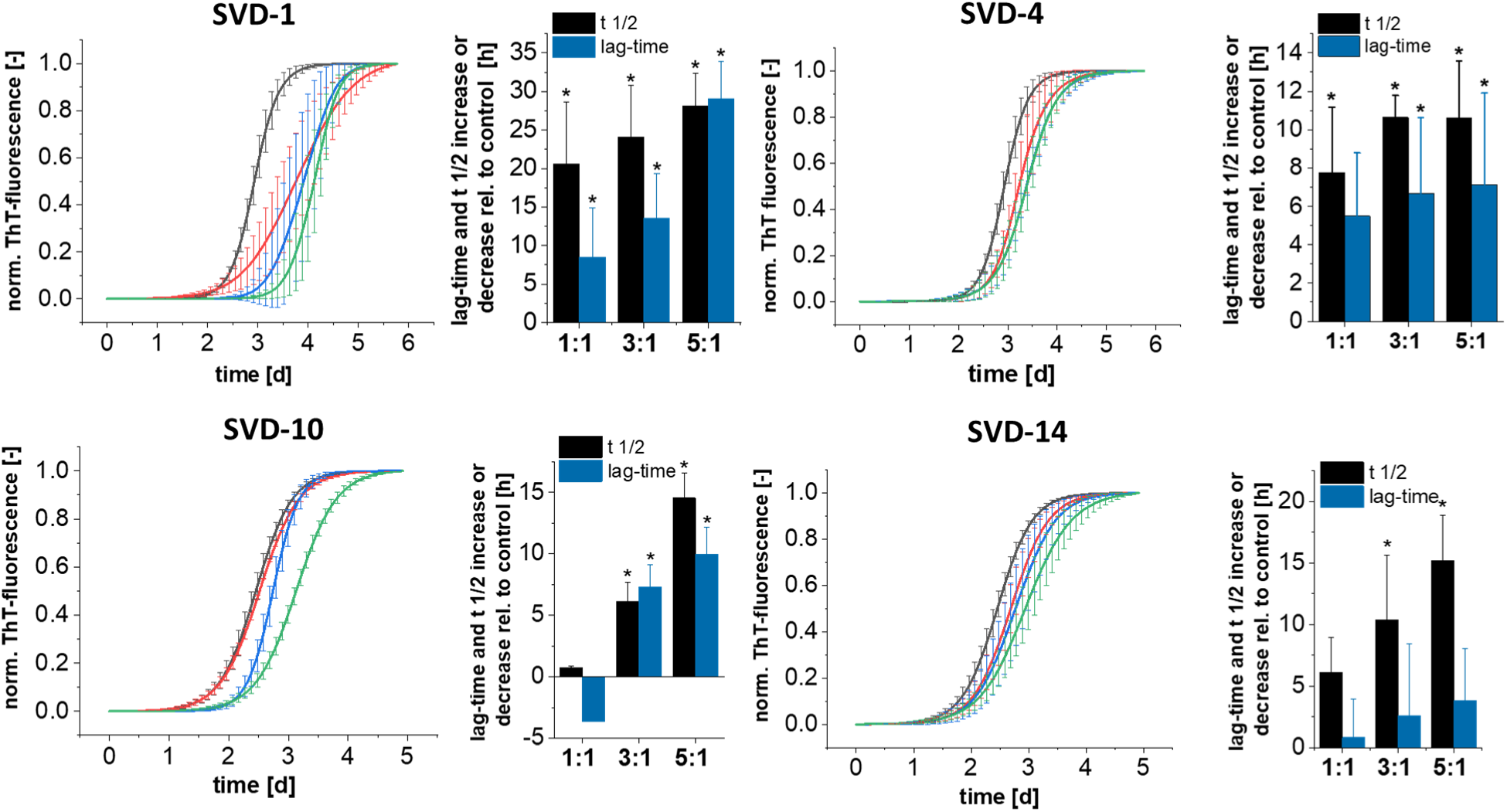
Thioflavin-T assay with α-syn and D-enantiomeric 16-mer peptides. 50 µM α-syn was incubated with or without 50, 150 or 250 µM D-peptide in PBS pH 7.4 (black: 0 µM; red: 50 µM; blue: 150 µM; green: 250 µM). ThT fluorescence progression was measured in a 96-well non-binding half-area plate (Corning, USA) with a Fluorostar platereader (BMG labtech, GE) at λex = 448 nm and λem = 482 nm. Samples were shaken at 300 rpm for 30 s per cycle using orbital shaking mode. The five replicates of each condition were individually fitted with a Boltzmann fit. The t ½ is given by the inflection point of the fit and the lag-time was approximated using the formula [lag-time = t ½ - 2dx] where dx is defined as the slope of the fit at x = t ½. t ½ and lag-time increase or decrease were determined based on the differences between samples with and without peptide, whereas the 0-value the reference control without inhibitor. The significance of time-shifts was tested with a two sample Welch‘s t-test with p < 0.05 (significant: *). Mean data shown with ± SD (n = 5). For α-syn without peptide the following t ½- and lag-times were determined: For SVD-1 and SVD-4: t ½: 70.4 h and lag-time 58.9 h for SVD-10 and SVD-14: t ½: 59.8 h and lag-time 49.4 h. All of the tested D-peptides decelerated α-syn aggregation, with SVD-1 being most efficient, with a t ½-time shift of + 28 h ± 4.2 h and a lag-time shift of + 29 h ± 4.9 h for a five-fold molar excess. Because of these results, we decided to continue with SVD-1 for further verification of the sequence specificity of the observed effects (Fig. 2).

**SI Figure 7:**
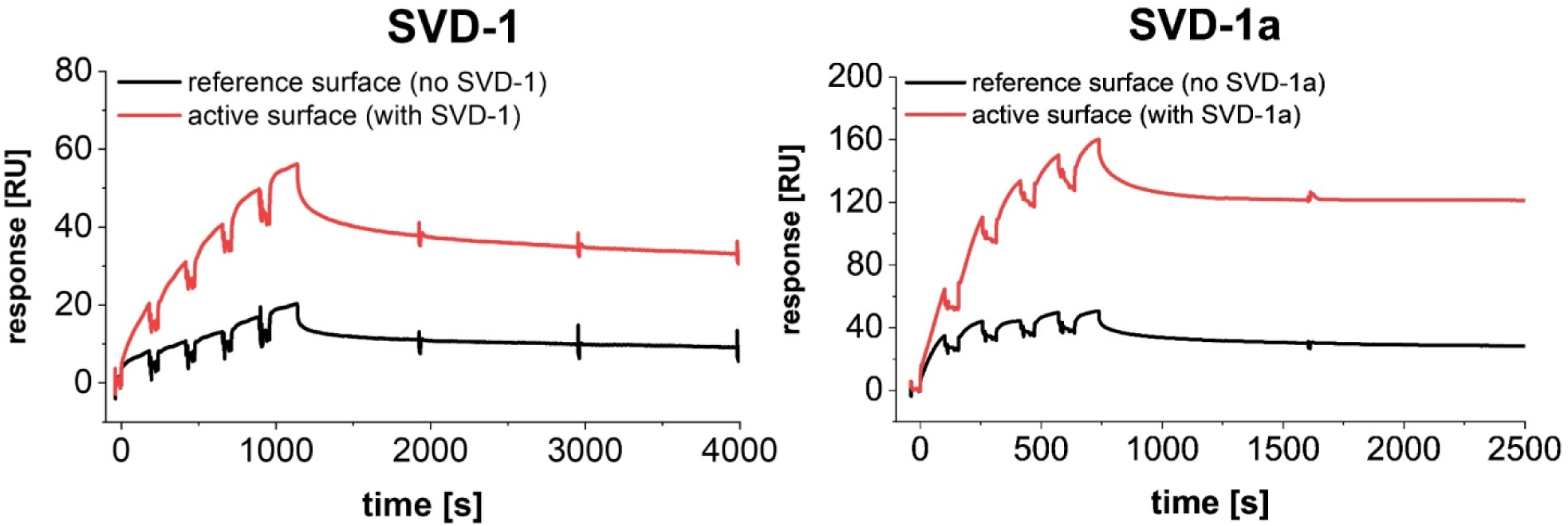
Single cycle injections of α-syn on active surface with immobilized SVD-1 and SVD-1a and the non-immobilized reference surface. SVD-1 (left) and SVD-1a (right) were immobilized on a carboxyldextran matrix via amino coupling until saturation was reached (CMD200M, Xantec, GE). Full-length a-syn was injected for 100 s for each concentration at 30 µl/min in PBS 7.4 on the active sensor with immobilized SVD peptide (red) and on the empty reference surface (black). Injections were performed using a serial dilution in the range of 30 to 500 nM and 30 to 150 nM for SVD-1 and SVD-1a, respectively. Referenced data are shown in Fig. 7.

**SI Figure 8:**
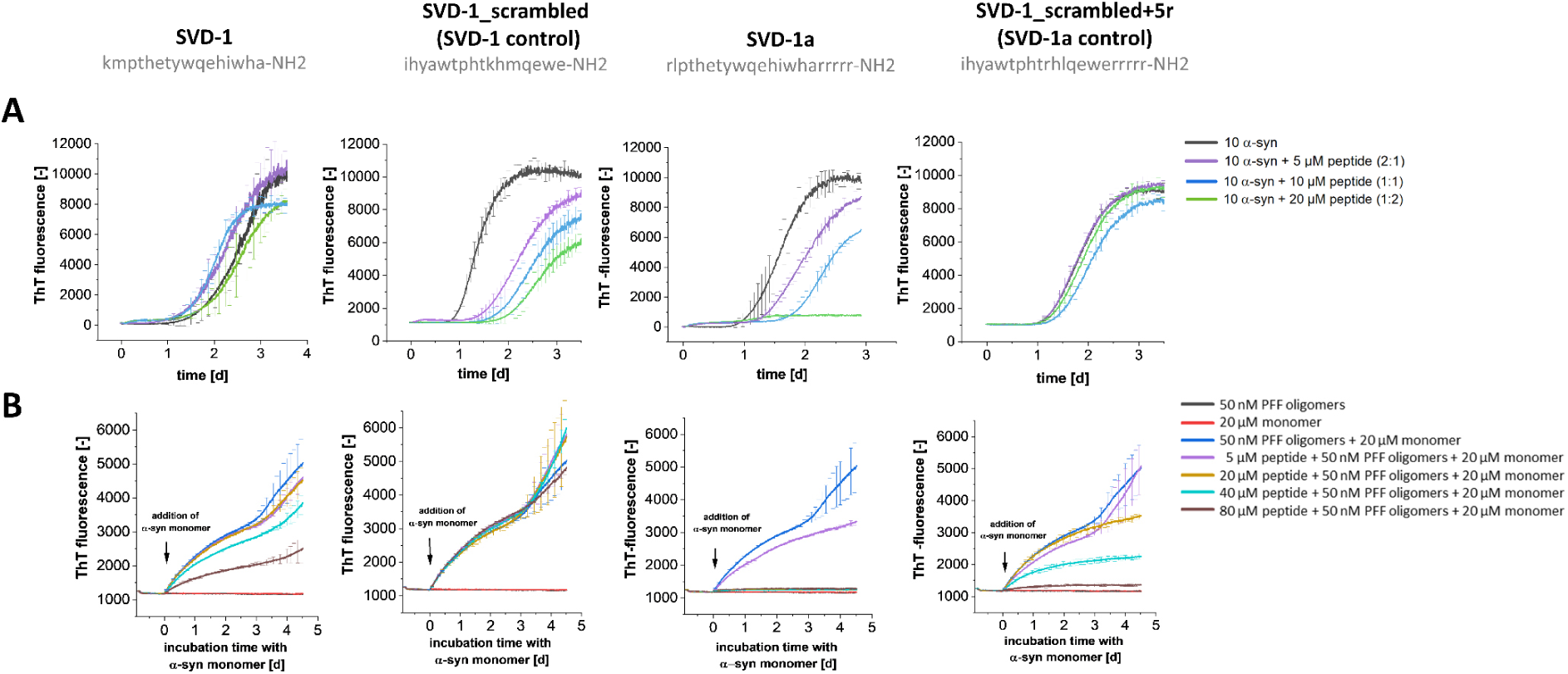
De novo and seeded ThT aggregation with SVD-1 and SVD-1a and the respective randomized control peptides (SVD-1_scrambled and SVD-1_scrambled+5r). ThT fluorescence progression was measured in a 96-well non-binding half-area plate (Corning, USA) with a Fluorostar platereader (BMG labtech, GE) at λex = 448 nm and λem = 482 nm. D-peptide sequences are shown as single letter amino acid code in grey. (**A**) *De novo* aggregation assay. 10 µM α-syn monomer was incubated with 5, 10 and 20 µM peptide at 37 °C, adding one borosilicate glass bead per well (d = 3.0 mm, Hilgenberg, GE) with continuous orbital shaking at 300 rpm in between reads. Mean data shown with ± SD (n = 5). The statistical evaluation on significance of inhibitory effects is shown in SI Fig. 10. Please note, when comparing the lag-times in this figure with those in SI figures 4 to 6, that shaking conditions were different. (**B**) Seeded aggregation assay. 50 nM monomer equivalent PFF oligomers as seeds were pre-incubated for 20 h with or without peptide at 37 °C under quiescent conditions. Only then, 20 µM α-syn monomer was added for induction of seeding and the incubation time with α-syn monomers started. Mean data shown with ± SD (n = 3).

**SI Figure 9:**
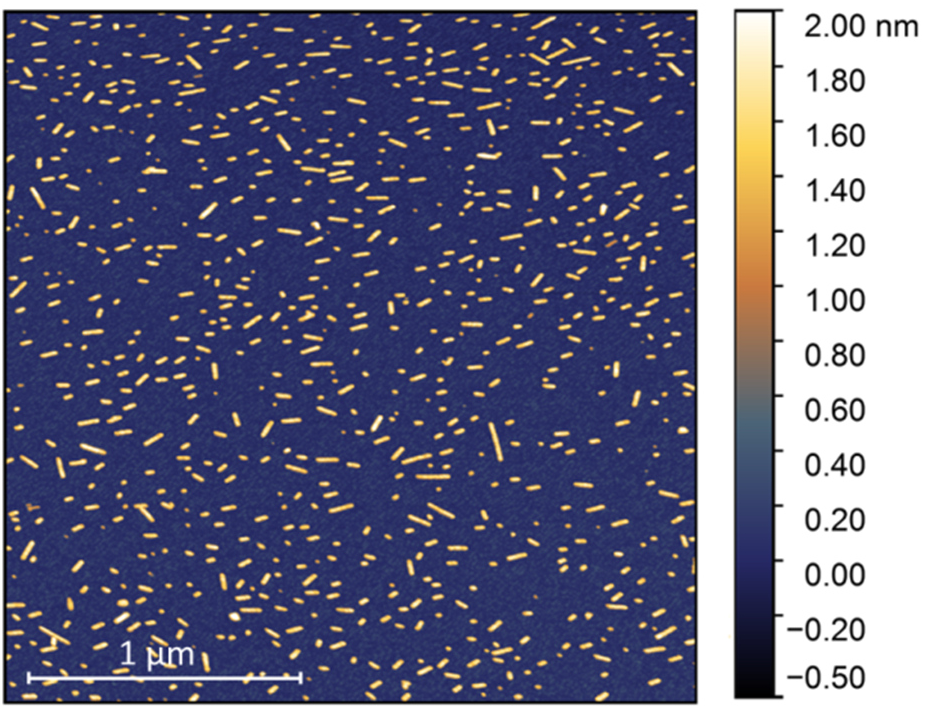
Atomic force microscopy image of α-syn PFF oligomer seed preparations. The preparation after ultracentrifugation was diluted to a final concentrations of 5 µM monomer equivalent in PBS pH 7.4. 5 µl was incubated for 30 min at RT on a freshly cleaved mica, washed three times with distilled water and dried in a gentle stream of N_2_. Measurements were performed in a Nanowizard 3 system (JPK Instruments AG, GE) using intermittent contact mode with 2.5 x 2.5 µm section and line rates of 0.5–1 Hz in ambient conditions with a silicon cantilever with nominal spring constant of 26 newtons/m and average tip radius of 7 nm (Olympus OMCL-AC160TS).

**SI Figure 10:**
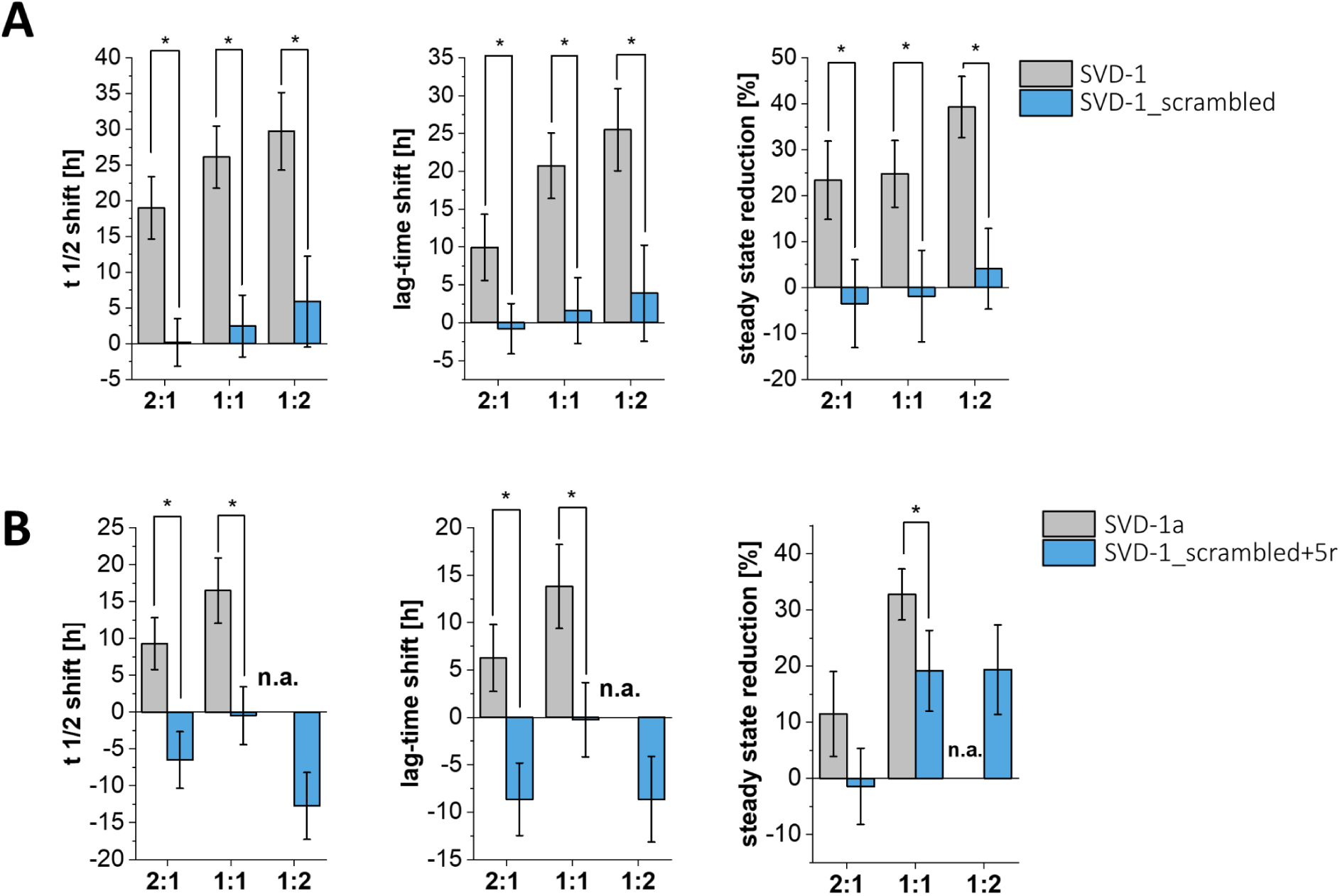
Statistical evaluation of *de novo* aggregation t ½-, lag-time shifts as well as steady state reduction for SVD-1 and SVD-1_scrambled as shown in SI Fig. 9. The five replicates of each condition were individually fitted with a symmetric Boltzmann fit. For each fit, x ½, lag-time and steady state response was determined in order to calculate their mean value for each condition. The t ½ is given by the inflection point of the fit and the lag-time was approximated using the formula [lag-time = t ½ - 2dx] where dx is defined as the slope of the fit at x = t ½. t ½ and lag-time shifts were determined based on the differences between samples with and without peptide, whereas the 0-value represents the reference control without inhibitor. The 2:1 stoichiometric sample of SVD-1a was not fitted due to lacking signal intensity and thus is marked with n.a.. α-Syn untreated control aggregation time: SVD-1: t ½ 32.9 h, lag-time 23.0 h; SVD-1_scrambled: 43.0 h; lag-time: 31.2 h; SVD-1a: t ½ 38.2 h; lag-time 27.2 h; SVD-1_scrambled+5r: t ½ 60.8 h; lag-time: 44.5 h. The significance of time-shifts between sample and the same concentration of the according control peptide was tested with a two sample Welch‘s t-test with p < 0.05 (significant: *). Mean data shown with ± SD (n = 5).

**SI Figure 11:**
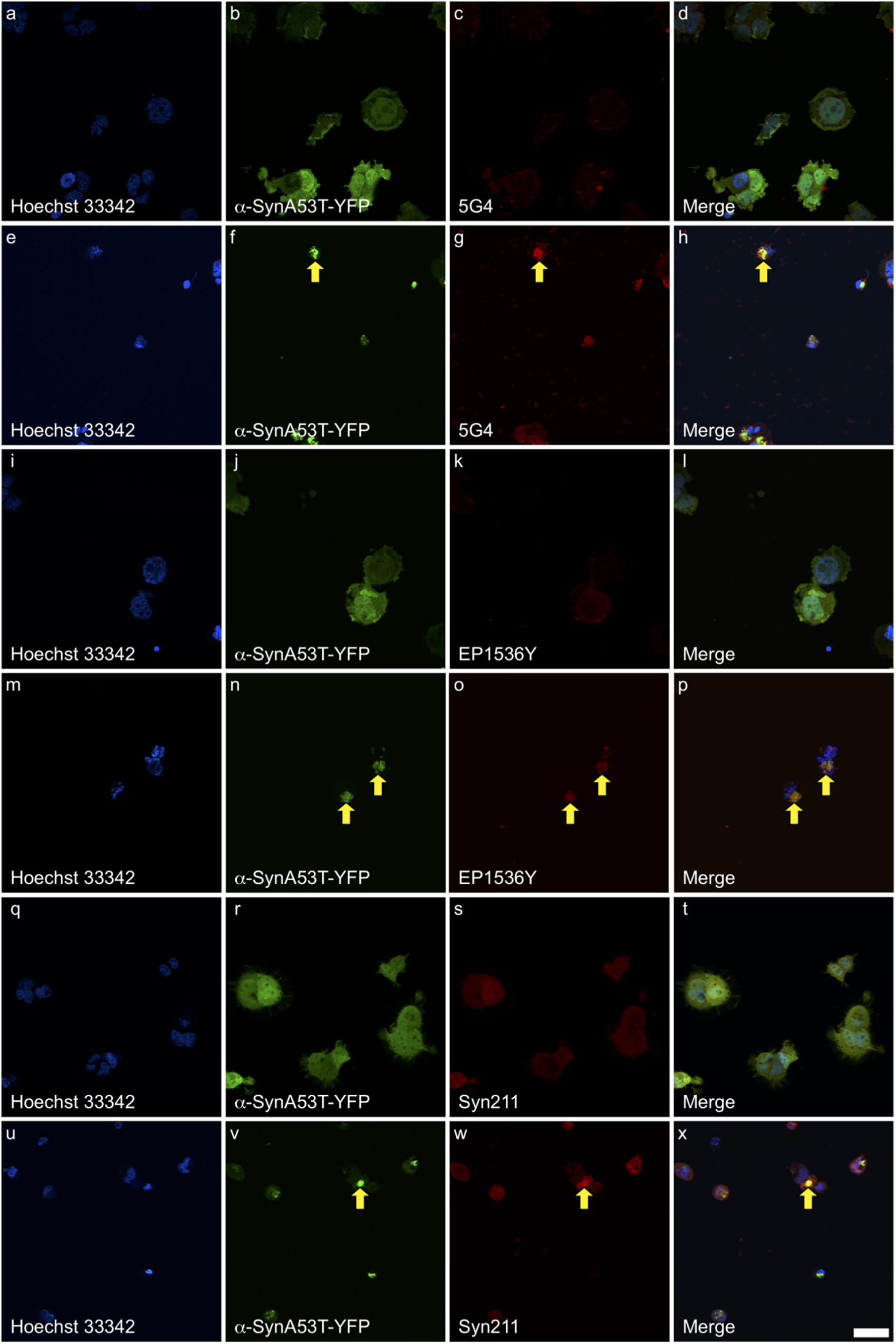
Immuofluorescence staining of α-syn aggregates in α-synA53T–YFP cells. After culturing unseeded cells (a–d, i–l, and q–t) and cells seeded with α-syn PFF oligomers s (e–h, m–p, and u–x) for 3 days, cells were fixed, permeabilized, and stained with fluorescently labeled antibodies (red). For detecting oligomeric and fibrillar α-syn, we used the anti-aggregated α-syn antibody, clone 5G4 (c, d, g, h). For detecting α-syn phosphorylated at serine 129, which accumulates when α-syn aggregates and forms deposits, we used the recombinant anti-α-syn (phospho S129) antibody EP1536Y (k, l, o, p). For detecting total αsyn, we used the anti-α-syn antibody syn211 (s, t, w, x). Nuclei were stained with Hoechst 33342. α-synA53T–YFP fluorescence is shown in green. Panels on the right represent merged images (d, h, l, p, t, x). While α-synA53T–YFP fluorescence is diffusely distributed in unseeded cells, it forms foci in cells seeded with α-syn PFF oligomers , indicative of α-syn aggregation. Yellow arrows indicate aggregated α-syn (yellow), a colocalization of antibody staining for α-syn (red) and fluorescence of aggregated α-synA53T–YFP (green). The scale bar indicates 100 µm.

**SI Figure 12:**
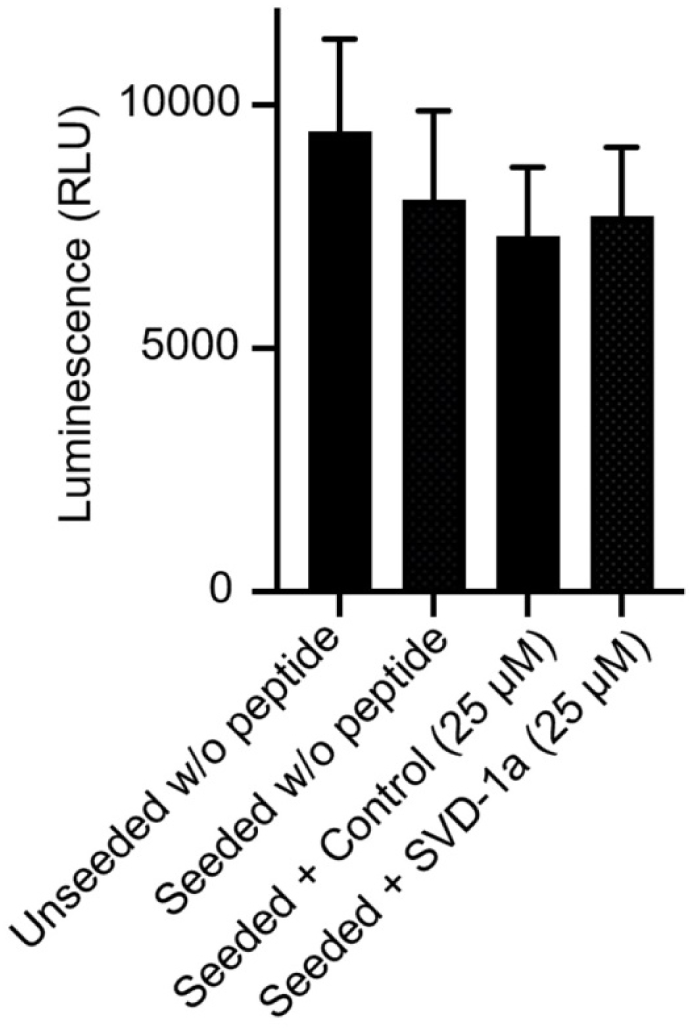
The transfection of PFF oligomers and SVD-1a does not significantly reduce HEK293 cell viability. CellTiter-Glo Luminescent Cell Viability Assay (Promega GmbH, GE) was used to determine the number of viable cells in culture based on quantitation of the ATP present, an indicator of metabolically active cells. After culturing cells in 384-well plates for three days, 35 µl of medium was removed from the wells and 40 µl of CellTiter-Glo Reagent directly added to each well. After mixing, luminescence was measured 10 min later using a Fluostar (BMG labtech, GE). The luminescence of seeded αSynA53T–YFP cells was slightly lower than that of unseeded cells, however, the difference was not significant. Also, treatment with a negative-control peptide or with SVD-1a did not induce significantly different luminescence in comparison to unseeded cells or seeded cells without peptide treatment. Significance was calculated using one-way ANOVA followed by Dunnett’s multiple comparisons test with p < 0.05.

**SI Figure 13:**
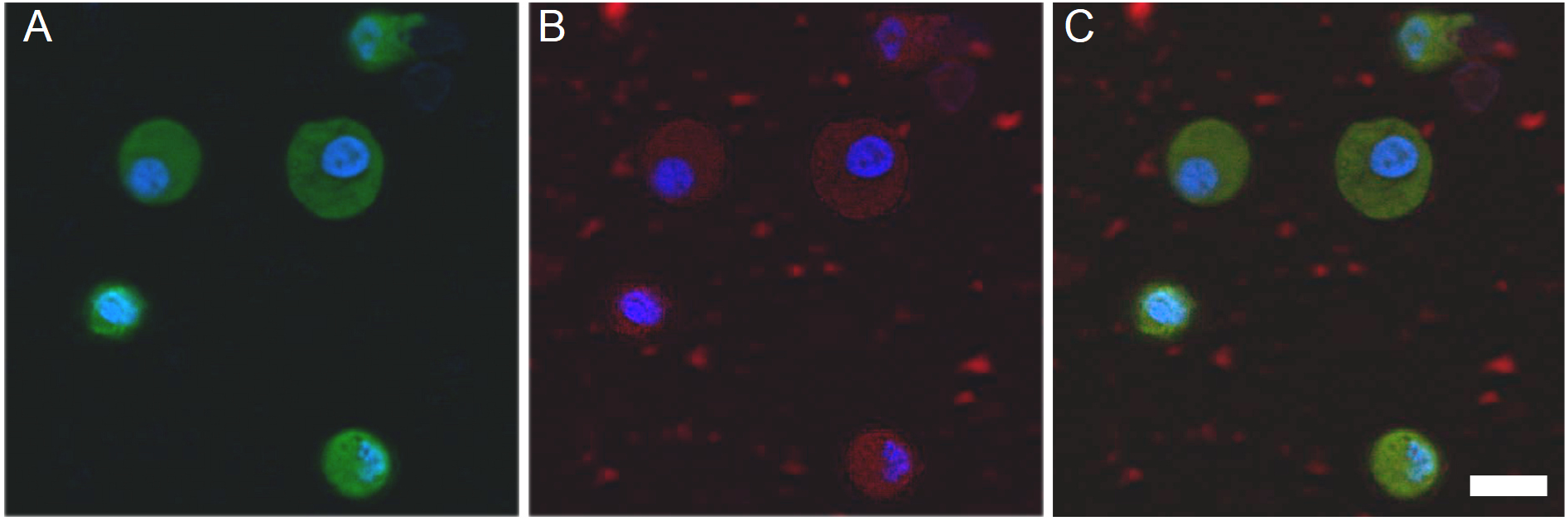
Fluorescently labeled SVD-1aCys_Alexa647 is taken up in α-synA53T–YFP cells. Unseeded α-synA53T–YFP cells were supplemented with 5 µM of SVD-1aCys_Alexa647 in the cell culture medium for 24 h. Confocal images of the cells show a diffuse green signal for cytoplasmic α-synA53T–YFP (**A**), and a diffuse red signal for SVD-1aCys_Alexa647 that was labeled with a red fluorescent dye and taken up by the cells (**B**). Nuclei were stained with Hoechst 33342 in blue. The last panel represents a merged image (**C**). The scale bar indicates 20 µm. Because the red fluorophore signal was homogeneously distributed in the cytosol of the cells, we assume that the peptide is able to enter the cells even without transfection.

**SI Figure 14:**
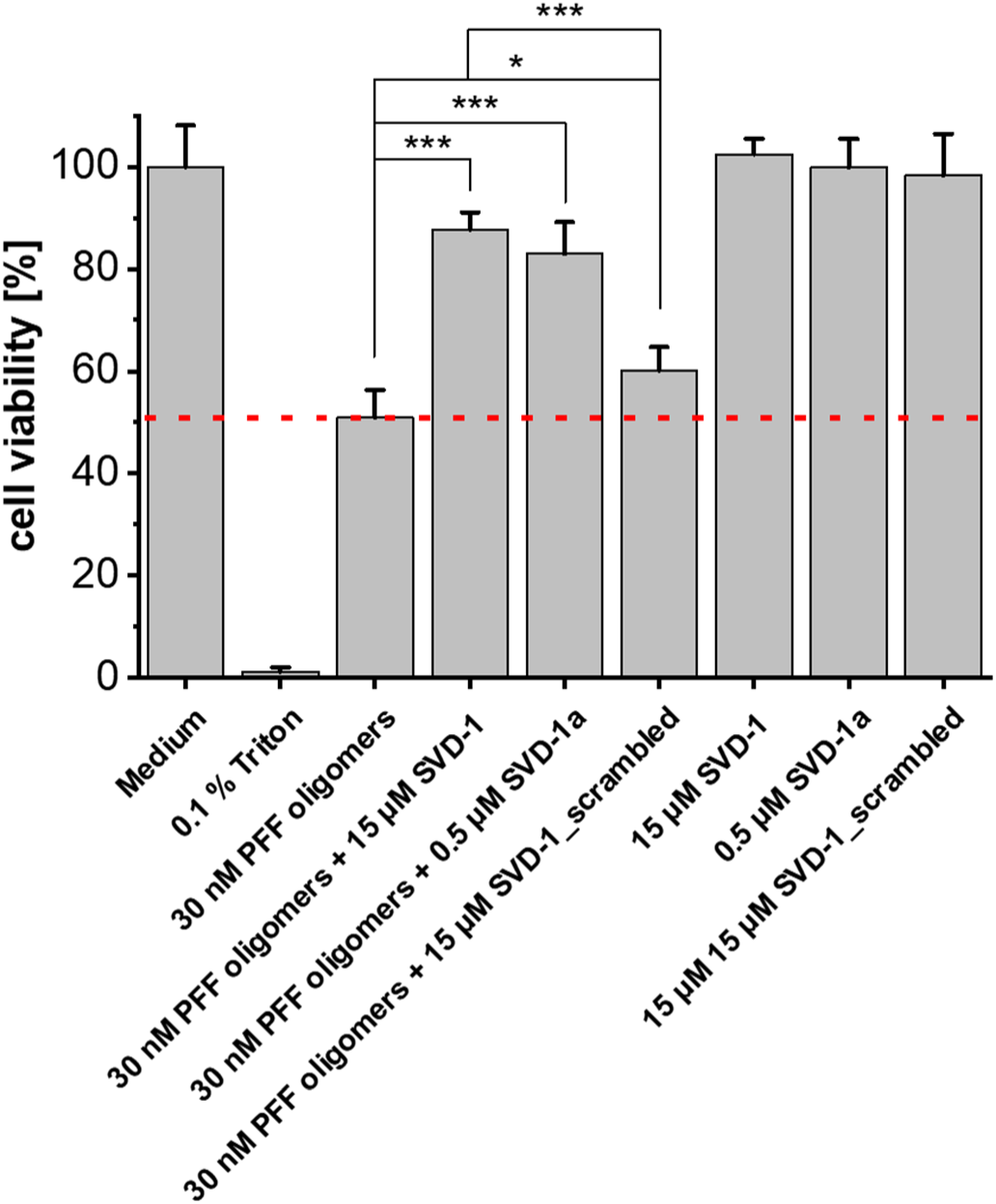
SVD-1 and SVD-1a increase cell viability in MTT assay with PFF oligomers seeds. SVD-1, SVD-1a SVD-1_scrambled were incubated over night at 37 °C and 300 rpm with or without 30 nM PFF oligomers and subsequently incubated with PC12 cells in final concentrations of 15 µM (SVD-1 and SVD-1_scrambled) or 0.5 µM (SVD-1a). While the cell viability was reduced with 30 nM PFF oligomers alone to 50 %, 89 and 82 % of cells were rescued in the presence of SVD-1 and SVD-1a, respectively. Slight increase of cell viability saving effect was also detected for 15 µM of the control peptide SVD-1_scrambled. Test on significance was performed using one-way ANOVA with Bonferroni post hoc analysis: * p ≤ 0.5, *** p ≤ 0.01. Mean data are shown with ±SD (n = 4). While the cell viability was reduced by 50 % in the presence of 30 nM soluble α-syn PFF oligomers alone, no toxic effect of the compounds on PC12 was observed. However, when the seeds were pre-incubated with SVD-1 or SVD-1a, the cell viability increased to 89 and 82 % using 15 µM or 0.5 µM compound concentrations, respectively. Thus, similar cell viability saving effects were found for SVD-1a using a 30-fold lower concentration. For the concentration of 15 µM SVD 1_scrambled, only a slight cell viability saving effect was found which was significantly lower than the cell saving effect identified for the same concentration of SVD-1. This result shows that the efficacy of the compounds on the soluble α-syn PFF is not only limited to a seeding capacity reduction, but additionally, the compounds also reduce the toxicity of the seeds in the cellular environment.

**SI Figure 15:**
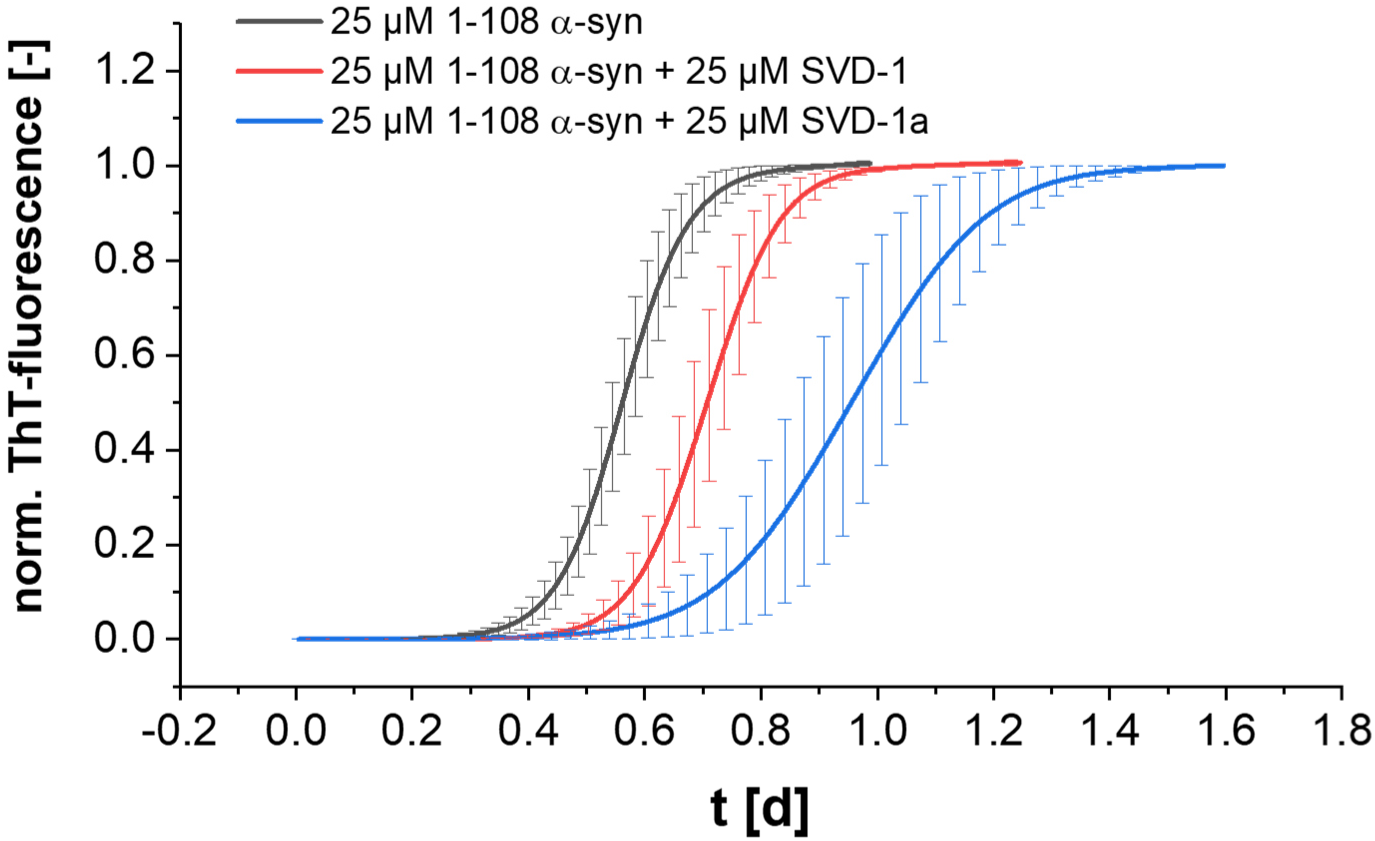
SVD-1 and SVD-1a inhibit the aggregation of 1-108 truncated α-syn. 25 µM monomeric 1-108 α syn was incubated with or without equimolar concentrations of SVD-1 and SVD-1a as described in the method section for *de novo* ThT-aggregation screening set-up. Aggregation curves were fitted with a symmetric Boltzmann fit and the steady state signal was normalized to 1. SVD-1 as well as SVD-1a show delay of 1-108 truncated α-syn aggregation.

